# Stochastic modelling of single-cell gene expression adaptation reveals non-genomic contribution to evolution of tumor subclones

**DOI:** 10.1101/2024.04.17.588869

**Authors:** M.G. Hirsch, Soumitra Pal, Farid Rashidi Mehrabadi, Salem Malikic, Charli Gruen, Antonella Sassano, Eva Pérez-Guijarro, Glenn Merlino, Cenk Sahinalp, Erin K. Molloy, Chi-Ping Day, Teresa M. Przytycka

**Affiliations:** National Library of Medicine, NIH, Bethesda, Maryland, USA; Department of Computer Science, University of Maryland, College Park, Maryland USA; University of Maryland Institute for Advanced Computer Studies, College Park, Maryland USA; Neurobiology Neurodegeneration and Repair Lab, National Eye Institute, NIH, Bethesda, Maryland, USA; Cancer Data Science Laboratory, Center for Cancer Research, National Cancer institute, NIH, Bethesda, Maryland, USA; Laboratory of Human Carcinogenesis, Center for Cancer Research, National Cancer Institute, NIH, Bethesda, Maryland, USA; Laboratory of Cancer Biology and Genetics, Center for Cancer Research, National Cancer Institute, NIH, Bethesda, Maryland, USA; Instituto de Investigaciones Biomédicas Sols-Morreale, Consejo Superior de Investigaciones Científicas, Universidad Autónoma de Madrid (IIBM, CSIC-UAM), Madrid, Spain

**Keywords:** clonal evolution, adaptation, Ornstein-Uhlenbeck processes, gene expression, single-cell analysis, melanoma, immunotherapy response, WNT signaling

## Abstract

Cancer progression is an evolutionary process driven by the selection of cells adapted to gain growth advantage. We present the first formal study on the adaptation of gene expression in subclonal evolution. We model evolutionary changes in gene expression as stochastic Ornstein–Uhlenbeck processes, jointly leveraging the evolutionary history of subclones and single-cell expression data. Applying our model to sublines derived from single cells of a mouse melanoma revealed that sublines with distinct phenotypes are underlined by different patterns of gene expression adaptation, indicating non-genetic mechanisms of cancer evolution. Interestingly, sublines previously observed to be resistant to anti-CTLA-4 treatment showed adaptive expression of genes related to invasion and non-canonical Wnt signaling, whereas sublines that responded to treatment showed adaptive expression of genes related to proliferation and canonical Wnt signaling. Our results suggest that clonal phenotypes emerge as the result of specific adaptivity patterns of gene expression.

## 1 Introduction

Cancer refers to a spectrum of diseases in which cells acquire the ability to evade normal mechanisms that control cell growth, differentiation, and movement. It is generally assumed that cancer originates from a single cell (or a small number of cells) and progresses through a branched evolutionary process, during which cells can acquire new mutations, undergo epigenetic reprogramming, and differentiate into distinct heterogeneous subpopulations, referred to as subclones. This clonal evolutionary process is generally assumed to be driven by the selection of cells exhibiting properties related to growth advantage like immunoevasion, evasion of cell death, and cell growth ^1,2,3^. Understanding these selective forces is important, yet studies investigating them have been limited and mutation-centric^4,5^. While mutations are fundamental to many aspects of cancer studies, their impact on cell function is often indirect, and cancer evolution also proceeds by epigenetic changes ^6,7,8^. A more direct way to uncover selective trends in tumor evolution due to the combined effect of genetic and non-genetic factors is to study adaptive changes in gene expression, because changes in gene expression mostly cause direct alteration of cell functions. Studying the evolutionary pressures acting on gene expression in light of clonal evolution can provide critical insights into the molecular changes that confer fitness advantage. Molecular changes caused by gene expression adaptation also impact response to specific therapies that modulate the gene expression, and thus the functions, of cells. Knowledge of evolution of gene expression can then enable the design of more effective treatment protocols.

Until recently, the ability to investigate such selective processes has been limited because of the use of bulk sequencing, which cannot differentiate between cell-level selection and and expression changes due to changes in cell populations. Single-cell RNA sequencing, on the other hand, gives us the opportunity to study gene expression evolution both within and across subclones, which otherwise is obfuscated when using bulk data. Single-cell RNA sequencing has been increasingly utilized to study dynamical changes in cell populations. Methods taking advantage of single-cell RNA sequencing include pseudotime analysis methods, which order cells along a trajectory based on similarities in their expression patterns (reviewed in^9^) and RNA velocity approaches, which employ splicing information to infer directed cellular changes ^10^. Single cell data has also been used with Ornstein-Uhlenbeck (OU) processes in order to investigate gene expression changes throughout cellular differentiation^11^. These methods are key in understanding the cellular differentiation process, but they do not investigate changes at extended evolutionary scales.

Other methods that examine a longer time frame use gene expression to identify clusters of tumor cells that comprise different subclones. They then perform differential expression analysis between the clusters to identify genes that vary between the subclones ^12,13,14^. However, not only are relationships between such clusters and tumor subclones often complex ^15^, but differential expression methods give no information about whether the difference in gene expression is due to selective pressures or neutral evolution over diversifying evolutionary distances ^16^.

In summary, no study has directly modeled evolutionary shifts on gene expression caused by selective pressures in the tumor microenvironment. Here, we perform the first formal study of such evolutionary changes of gene expression. Specifically, we leverage single-cell full-length RNA sequencing data to investigate the role of selection on gene expression and function in subclonal evolution of cancer. We model clonal gene expression evolution using stochastic Ornstein-Uhlenbeck (OU) processes which have been previously applied in the context of species evolution ^17,18,19,20,21,22,13^ and cellular differentiation ^11^, but have not been utilized in studies on subclonal evolution. Our use of OU processes allows us to model changes in subclone-specific expression along trajectories defined by the evolutionary history of the subclones. Utilizing the evolutionary history enables us to evaluate whether gene expression is shifting because it is subject to adaptive evolution as opposed to shifting due to random changes caused by neutral evolution. Further, using single-cell data allows us to distinguish between subclonal growth and adaptive evolution, which is critical to understanding selective pressures in cancer and is not possible with differential expression analysis, even when clusters accurately reflect subclones.

To evaluate the capabilities and limitations of the OU processes, we first evaluated the performance of the models on realistic simulations of single-cell RNA tumor evolution data. We then used the OU processes to model the evolution of the M4 melanoma^1^, which exhibited intratumoral heterogeneity^23^ (Figure 1). Specifically, we applied these stochastic processes on multiple “clonal sublines” of an M4 melanoma cell line (known as B2905)^2^ (Figure 1a,b). In our analysis, we incorporated knowledge of the evolutionary history of the sublines and their phenotypes, such as aggressiveness and response to immunotherapy (Figure 1c) and identified genes with adaptive expression related to these phenotypes (Figure 1d). We found that different phenotypes were associated with adaptation of gene expression for distinct genes and pathways. Included in these pathways were alternative Wnt signaling pathways, implying an association between clonal selection for alternative Wnt signaling and immune response (Figure 1e). By subjecting the parental M4 melanoma to anti-CTLA-4 treatment, we further found that genes with adaptive expression in the treatment resistant sublines captured by our analysis display expression in post-treatment tumors consistent with immunotherapy response (Figure 1f).

**Figure 1:**
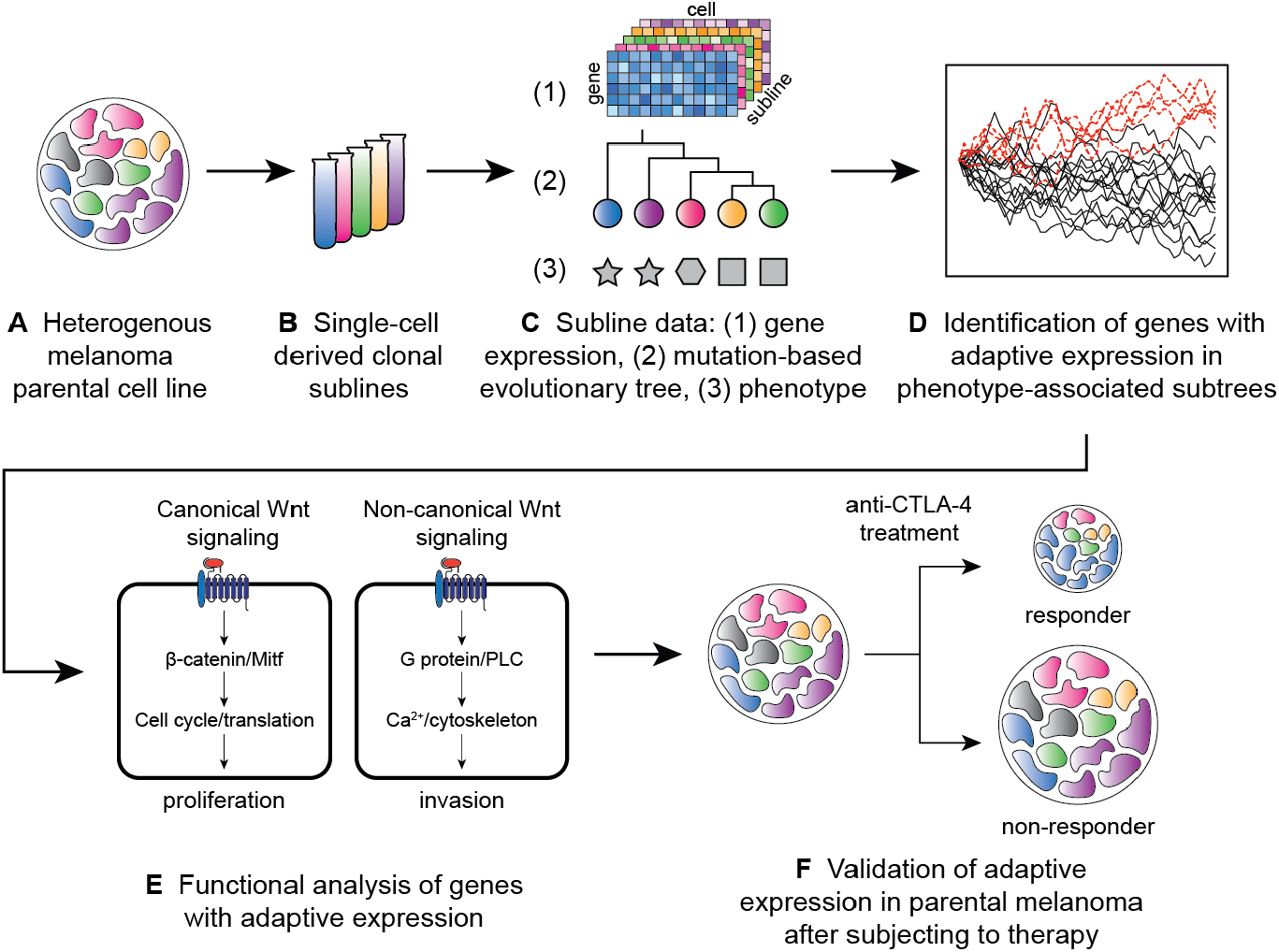
Summary of approach. A heterogeneous melanoma cell line **(A)** is used to obtain single cell-derived sublines **(B)**. Single-cell gene expression data, a mutation-based phylogenetic tree, and phenotypic features of the tumors derived from the sublines **(C)** are used to identify genes with adaptive expression associated with the cancer phenotypes **(D)**. The genes with adaptive expression are subjected to functional analysis **(E)**. For additional validation, tumors seeded from the parental line are subjected to anti-CTLA treatment. Overlap between genes with adaptive expression and genes differentially expressed between post-treatment responder and non-responder tumors are compared **(F)**.

Our study provides evidence that evolution of distinct phenotypes in cancer is driven by selection of subclones based on gene expression. We present novel use of single-cell RNA sequencing, a mutation-based phylogeny, and phenotypic information for evaluating evolutionary selective forces acting on gene expression. Such analysis could inform targeted treatment for tumors exhibiting different phenotypes to increase the efficacy of the treatment and decrease the chance of tumor recurrence.

## 2 Results

### 2.1 Gene expression adaptation of clonal evolution in cancer can be modeled with Ornstein-Uhlenbeck processes applied to sc-RNA data

#### Experimental Setting

To study gene expression evolution in cancer, we analyze data that was previously generated in Gruen et al. ^24^; here we briefly review the data generation protocol. When implanted in syngeneic C57BL/6 mice, B2905 melanoma cells form tumors that exhibit heterogeneous responses to immune checkpoint blockade (ICB) therapies among individual hosts ^23^. To reconstruct the subclonal evolutionary history of the melanoma, 24 single cells were first isolated from the parental B2905 cell line and expanded to become individual clonal sublines, C1 to C24 (Figure 2a, box 1), as described previously ^24^. The sublines were then subjected to bulk whole-exome sequencing (Figure 2a, box 3). Mutations were called from the exome data of the sublines and used to build a mutation-based phylogeny, in which the sublines represent subclones derived from the melanoma cell of common origin ^15^. Next, for each subline, eight cells were isolated for full-length single-cell RNA sequencing (Figure 2a, box 2). Cells were filtered based on quality control measures, leaving one to eight gene expression measurement replicates per subline. Mutations were called from the single-cell RNA data and also used to build a phylogeny of the sublines. A consensus tree was constructed between the DNA and RNA mutation-based phylogenies as described in Mehrabadi et al. ^15^, and this tree is what is used in this study (Figure 2b)^3^. Cells from the individual sublines were then implanted into distinct but genetically identical mice and in vivo growth information was recorded (Figure 2, box 4).

**Figure 2:**
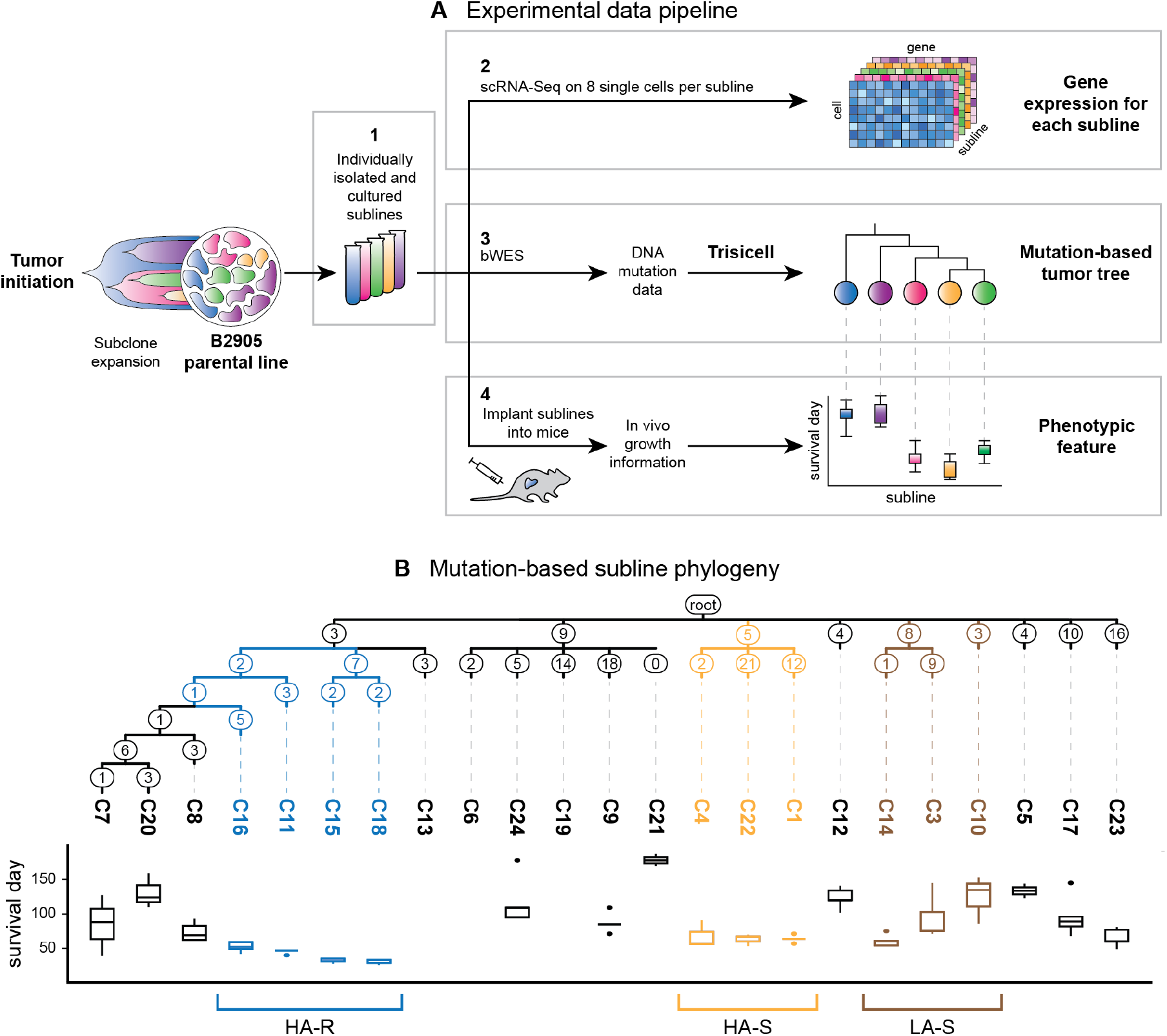
**(a)** Schematic illustrating the generation of the clonal sublines from a mouse melanoma cell line. Box 1: 24 single cells are isolated from the parental B2905 cell line and cultured into sublines. (Only five shown for illustration purposes.) Box 2: The sublines undergo single-cell RNA sequencing, resulting in gene expression data for each subline. Box 3: The sublines undergo bulk whole-exome sequencing and mutation calling. The mutation data is used to create a mutation-based tumor tree using Trisicell^15^. Box 4: The individual sublines are implanted into mice and allowed to grow. In vivo growth information for each subline-derived tumor is recorded, resulting in distinct phenotypes (time to reach a pre-defined size) across sublines. These three resulting data sources are used as input to the stochastic model. **(b)** Consensus mutation-based phylogeny tree estimated by Trisicell and growth kinetics of the clonal sublines ^15^. The branches are labeled with the number of expressed mutations along that branch, which are used as the branch lengths of the phylogeny. Sublines are colored based on their phenotypes: high aggression and resistant (HA-R) sublines (C16, C11, C15, C18) are blue, high aggression and sensitive (HA-S) sublines (C4, C22, C1) are orange, and low aggression and sensitive (LA-S) sublines (C14, C3, C10) are brown. Growth kinetics, measured in the time the tumor took to reach a predefined size, are shown below each subline. Sublines that did not reach the set tumor size in the time frame have no growth kinetics shown.

In their original analysis of the data, Mehrabadi et al. ^15^ found that sublines of major clades in the phylogeny exhibited distinct growth kinetics in vivo (measured by the survival time of the hosts following implantation of the cells to grow tumors; Figure 2b) and putative sensitivity to ICB treatment. We can divide the subline groups based on aggressiveness and treatment sensitivity. With respect to aggressiveness, the sublines can be divided into high aggression (HA) (fast growth rate) and low aggression (LA) (slow growth rate). In terms of treatment sensitivity, they can be divided into resistant (R) and sensitive (S) to ICB treatment. The following three diverse phenotype-driven subline groups were chosen for evaluation based on the phenotypes observed in the experimental data of Mehrabadi et al. ^15^: (1) sublines C11, C15, C18, and C16, which exhibit high aggression and are resistant to ICB (HA-R); (2) sublines C1, C4, and C22, which exhibit high aggression and are sensitive to ICB (HA-S); (3) sublines C3, C10, and C14, which exhibit low aggression and are sensitive to ICB (LA-S). The varying characteristics of these sampled sublines demonstrate a heterogeneity within the cell population of the parental line.

#### Model

To understand how gene expression in different sublines shifts due to selective pressures, we use stochastic models to estimate such shifts for each of the phenotypically distinct groups of sublines. Finding genes that adapt differently between the sublines will give insight into how the sublines have adapted to exhibit contrasting growth rates and resistance to ICB therapies.

Our approach to modeling clonal gene expression evolution builds upon Ornstein-Uhlenbeck (OU) processes that are often used to study the evolution of gene expression across species ^25,20,21,26,22,17^. Instead of species, we apply our method to the sublines evolved from a cell of common origin. We assume that the evolution of gene expression in a tumor is a stochastic process that follows one of three alternative evolutionary models: neutral (no selection), constrained (putative negative selection), or adaptive (expression shift consistent with clone(s) adaption) evolution. Neutral evolution is expected to occur when there are no selective pressures on the expression. We model neutral evolution with a Brownian motion (BM) process, where changes to gene expression occur randomly at each time step (Figure 3a, box 1). Constrained evolution is expected when there is a selective pressure against a divergence from a common optimal value acting on all sublines. We model constrained evolution with an OU process with a single optimum: the gene expression for all sublines converges towards a common value, *θ*_0_ (Figure 3a, box 2). Adaptive evolution is expected to occur when selective pressures cause the sublines to adapt (shift expression) to different values (Figure 3a, box 3). We model adaptive evolution as an OU model with two optima^4^, *θ*_0_ and *θ*_1_. In order to evaluate adaptive evolution, the sublines must be partitioned into groups expected to have different optima values; this partitioning is called a “regime.” In this setting, one set of the sublines is the group of interest, hereafter called the “chosen” sublines, and the rest of the sublines are considered as “background.” In species evolution, such adaption might be a result of differing environments. In clonal evolution, adaptation might be the result of differing locations inside the tumor, and thus varying pressures of the tumor microenvironment, and/or differing responses to the same selective pressures. Further details of BM and OU processes can be found in Methods Section 3.1.

**Figure 3:**
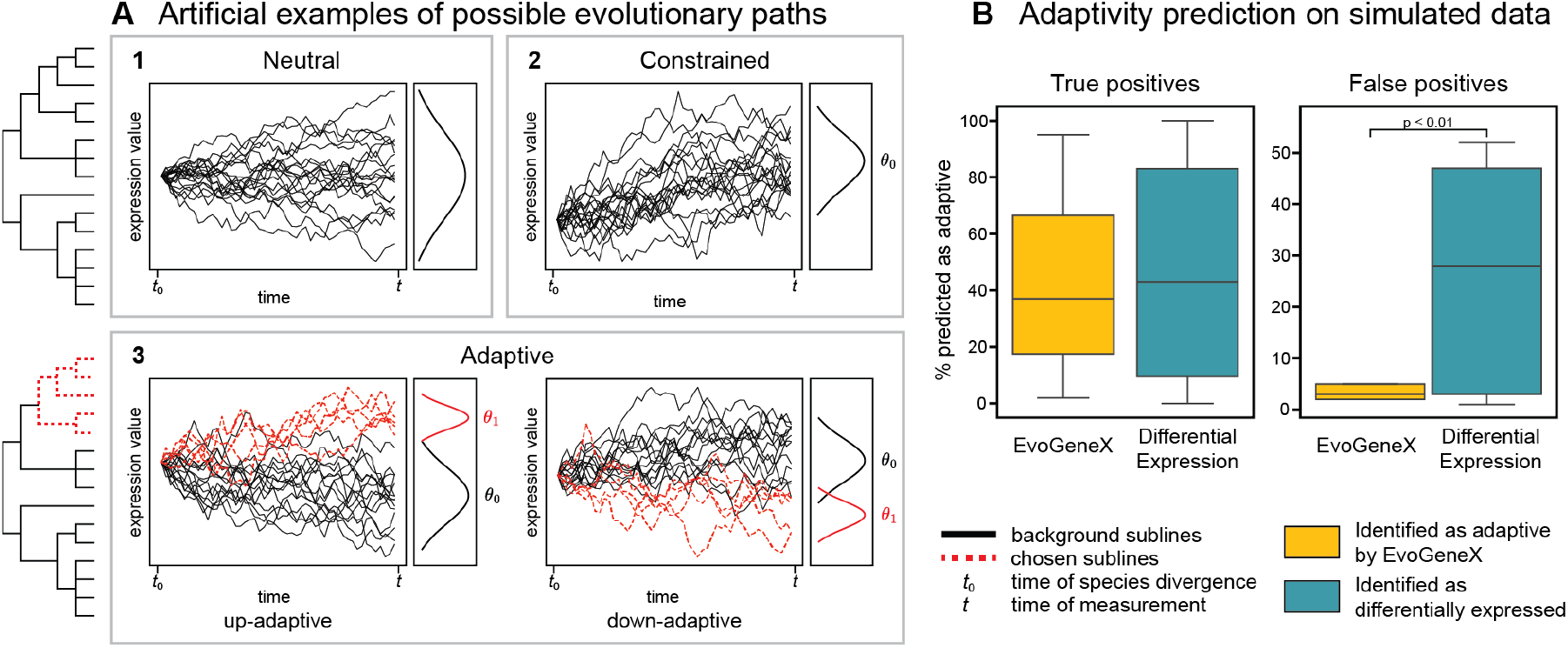
**(a)** Examples of possible evolutionary trajectories of gene expression along the phylogeny. On the left is the phylogeny that was used to simulate the data. For neutral (Box 1) and constrained (Box 2) evolution, no phenotypic feature is used and all sublines are considered to have the same dynamics (solid black branches). In adaptive evolution (Box 3), some sublines have different evolutionary dynamics (dashed red branches). Each box shows a simulated example evolutionary trajectory of expression for the different sublines as well as the resulting expression distribution that would be observed and used as input to the model. Optima values (*θ*) used to simulate the data are indicated where applicable. **(b)** Adaptivity and differential expression predictions. Right: Percentage of adaptively simulated genes correctly identified to be adaptive or differentially expressed across all simulations. Left: Percentage of neutrally simulated genes incorrectly identified to be adaptive or differentially expressed across all simulations. Results from EvoGeneX are shown in yellow; results from differential expression are shown in teal.

To detect constrained and adaptive evolution of gene expression from single-cell RNA sequence data of the sublines, we use EvoGeneX, a recently introduced software ^17^. EvoGeneX takes as input the phylogenetic tree, the expression data for the gene under evaluation, and the regime for the adaptive evolution model; it outputs whether the gene can be considered to be constrained or adaptive. We evaluate three regimes of interest based on the groups of sublines sharing phenotypic properties described above: high aggression and resistant (HA-R), high aggression and sensitive (HA-S), and low aggression and sensitive (LA-S). The internal nodes are assigned optima based on the shared ancestry of the chosen sublines in that regime (see Figure 2b). We run EvoGeneX on each regime independently. For each regime, all sublines not in the group under evaluation are used as the background for comparison.

EvoGeneX detects adaptation for each gene as follows. First, it uses maximum likelihood to estimate the model parameters and calculate a likelihood value for each of the three models (BM, OU with one optimum, and OU with two optima). The stochastic processes are computed down the branches of the tree, with the number of steps of each process proportional to the branch lengths, and lineages sharing internal branches sharing the same values. Second, it calculates a p-value by using likelihood ratio tests to determine if neutral evolution can be rejected for constrained or adaptive evolution. After running EvoGeneX for all genes, we correct p-values using the Benjamini-Hochberg false discovery rate (BH-FDR) method. Genes are labelled as constrained or adaptive if the corrected p-value from the likelihood ratio test against neutral evolution is less than 0.05.

Because EvoGeneX estimates the parameter values of each model, the adaptive genes can be considered to be adaptively up-regulated or adaptively down-regulated by comparing the estimated optima values of the chosen sublines in the regime of interest and the background: if the chosen sublines have a higher estimated optimum than the background, the genes are considered to be adaptively up-regulated, and if they have a lower estimated optimum they are considered to be adaptively down-regulated.

EvoGeneX utilizates biological replicates to estimate within-species expression variability. The use of replicates is vital when using OU processes to reduce false positive results^22,27,17^. In the case of species, the biological replicates are different individuals of the same species. In our setting, the individual cells sequenced for each subline serve as these biological replicates. Accounting for such within-subline variation decreases potential false positives that could result from high variation within each subline as well as from the noise inherent in single cell data.

#### Applicability of the OU-based model for single cell cancer data and validation with simulations

Previous studies inferring gene expression evolution with Ornstein-Uhlenbeck (OU) models utilised bulk data from tissues, organs, or even whole organisms ^18,26,17^. However, bulk data represents the average expression across all the cell types in the population. Because of this aggregation, when the composition of the cellular population changes, the bulk expression value also changes. Such settings do not allow for differentiating expression shift driven by evolution within specific cell types from shift driven by evolution of tissue composition ^27^. Distinguishing evolutionary changes from compositional changes is particularity relevant in the context of cancer, where subclonal growth and selection change the cellular composition of the tumor. Bulk expression analysis is unable to distinguish gene expression shift due to differential subclonal growth from gene expression evolution within individual cells in the subclones. Therefore, in this study, we leverage single cell derived gene expression. However, single cell data comes with unique challenges, especially high noise. To confirm that OU-based modeling is effective for such data, we perform simulation studies.

In our simulation studies, we generated expression data using an OU process on the evolutionary tree inferred for our data (Figure 2b) ^15^ under different modes of evolution. For the constrained and adaptive simulations, we varied the parameters to reflect both strong and weak signals of adaptation. We added varying levels of noise to reflect the nature of single cell measurements, aiming to obtain similar expression patterns as in the experimental TPM data. (Details on simulated data generation can be found in Supplementary Section 1.1; the simulated data is provided in the Supplementary files.) We then evaluated whether our approach using EvoGeneX could correctly infer the evolutionary model used to generate the data. Our simulation studies confirmed that in this setting EvoGeneX can uncover expression adaption, with the accuracy depending on the model parameters: the distance between the optima, the pull towards the optima, and the within-subclone variation. Depending on these parameters, EvoGeneX correctly identified up to 95% of genes simulated to have adaptive expression as adaptive (identifying 42% across all parameter combinations). It also had a low false positive rate: of all neutrally evolving experiments, a total of only 3.4% genes were predicted to be adaptive (Figure 3b). However, in this setting EvoGeneX could not reliably distinguish constrained evolution from neutral evolution. Thus in the remainder of the paper we focus on inferring adaptation only. (Details of results can be found in Supplementary section 1.2; full results is provided in the Supplementary files.)

Finally, we used the simulation study to test if OU-based models provide an advantage over differential expression analysis as a basis to infer adaptation. We performed differential expression (using an independent t-test on the simluated TPM expression data) between the genes simulated in the chosen sublines and background sublines. In real data, the groups are unknown and clustering methods are used to identify them. We use the known divisions rather than a clustering step to better compare differential expression analysis and EvoGeneX.

More adaptive genes were detected to be differentially expressed than were identified by EvoGeneX to be adaptive (across all simulations 46% of genes are differentially expressed compared to 42% being labeled as adaptive by EvoGeneX), though the difference is not statistically significant (Figure 3b). However, differential expression had a much higher false positive rate of adaptive predictions: 26.3% of neutrally simulated genes were identified as differentially expressed compared to 3.4% identified as adaptive by EvoGeneX (statistically significant (p*<*0.01); (Figure 3b)). Although differential expression can sometimes identify more adaptive genes than are predicted by EvoGeneX, the high false positive rate of differential expression demonstrates that our method using EvoGeneX is applicable to single-cell RNA sequence data and is more useful for identifying potential adaptation in this setting than differential expression analysis alone. (Details of results can be found in Supplementary section 1.2; full results can be found in the Supplementary files.)

### 2.2 Gene expression adaptivity is distinctly constrastive in subclones with different phenotypes

Given the distinct subclonal phenotypes observed, our goal was to identify genes with expression levels under adaptive evolution driving the properties of each of the high aggression and resistant (HA-R), high aggression and sensitive (HA-S), and low aggression and sensitive (LA-S) subline groups. To address this question, we used EvoGeneX to identify genes with expression patterns consistent with adaptive evolution for each regime as described in Section 2.1.

Of the 8,363 genes analyzed, we identified 812 (9.7%) genes with adaptive expression in HA-R (496 adaptively up-regulated and 316 adaptively down-regulated), 1,277 (15.3%) genes with adaptive expression in HA-S (201 adaptively up-regulated and 1,076 adaptively down-regulated), and 616 (7.4%) genes with adaptive expression in LA-S (288 adaptively upregulated and 328 adaptively down-regulated). Consistent with our simulation studies, we found no constrained genes; see the Supplementary Section 2 for further discussion of this finding.

We performed KEGG enrichment analysis to investigate the functions of the genes with adaptive expression in each regime. Because we expect that the genes with adaptively upregulated and adaptively down-regulated expression will have different functions, we run enrichment analysis on them separately. The genes with adaptively down-regulated expression in HA-R and adaptively up-regulated expression in HA-S are enriched for “ribosome^5^.” The genes with adaptively down-regulated expression in HA-S are enriched for “Rap1 signaling pathway” and “bacterial invasion of epithelial cells.” The genes with adaptively up-regulated expression in LA-S are enriched for “pathways of neurodegeneration.” The pathway enrichment for each regime is shown in Figure 4a and the specific genes involved in each enrichment are listed in the Supplementary files. There is no significant enrichment in KEGG pathways for the genes with adaptively up-regulated expression in HA-R or adaptively down-regulated expression in LA-S.

**Figure 4:**
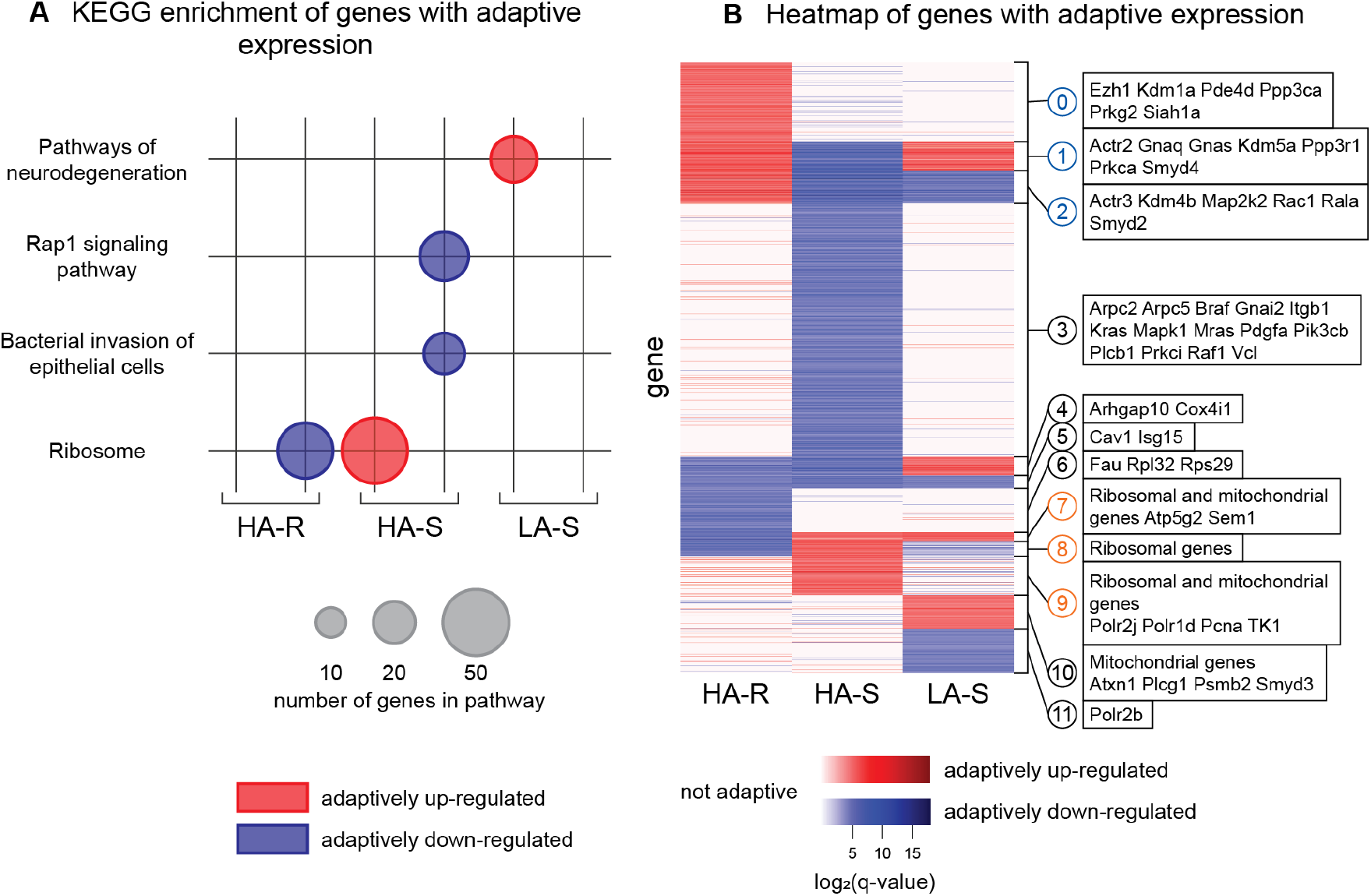
**(a)** KEGG enrichment of genes with adaptive expression, divided by regime and expression regulation direction. Genes with adaptively up-regulated expression for each subline group are shown in red on the left, genes with adaptively down-regulated expression are shown in blue on the right. The size of the dot represents the number of genes present in that term. **(b)** Heatmap of genes with adaptive expression, grouped into 12 clusters, and example genes in each cluster. Each row is a gene that has adaptive expression in at least one subline. Colors are based on the log of the q-value for each gene, with adaptively up-regulated expression shown in red, and adaptively down-regulated expression shown in blue. Genes with expression that is not adaptive in a subline group are shown in white. Clusters labeled in blue have adaptively up-regulated expression in high aggression and resistant (HA-R). Clusters labeled in orange have adaptively up-regulated expression in high aggression and sensitive (HA-S). See Methods section 3.1 for clustering methods.

We performed k-means clustering on the genes with adaptive gene expression in at least one of the regimes (see Methods Section 3.1 for details) and present the results in a heatmap (Figure 4b). As can be seen, there are large clusters of genes that are largely only adaptive in one regime (e.g. clusters 0, 3, and 6), but there is also distinct overlap between the regimes. For example, genes in clusters 1 and 2 have adaptively up-regulated expression in HA-R but adaptively down-regulated expression in HA-S, while genes in clusters 7 and 8 have the opposite adaptive expression pattern between these two regimes. In parallel, genes in clusters 1 and 4 have adaptively up-regulated expression in LA-S but adaptively downregulated expression in HA-S.

We then investigated how the genes associated with enriched pathways were distributed across clusters. Noticeably, clusters 1, 2, and 3, which are composed of genes with adaptively down-regulated expression in HA-S, contain the genes that are in the enriched pathways “Rap1 signaling” and “Bacterial invasion of epithelial cells.” Many of these genes also have adaptively up-regulated expression in HA-R (clusters 1 and 2). In contrast, clusters 7, 8, and 9, which are composed of genes with adaptively up-regulated expression in HA-S, contain the genes that are in the enriched “Ribosome” pathway, and many also have adaptively downregulated expression in the HA-R sublines (clusters 7 and 8). Together, these results suggest that the sublines with contrasting phenotypes evolved to have distinct transcriptional states.

### 2.3 Functional implications of genes with the adaptive expression pattern in melanoma

In this section, we provide more detailed biological interpretation of the adaptivity profiles between the sublines. Specifically, we analyze the genes within the clusters with adaptively upregulated expression in each of the high aggression and resistant (HA-R) and high aggression and sensitive (HA-S) subline groups.

We first consider the genes with adaptively up-regulated expression in HA-R (clusters 0, 1, and 2). We found many genes involved in G protein-coupled receptor (GPCR) signaling and cell invasion. They include G protein signaling (*Gnaq, Gnas*, and *Rab5b*), their direct downstream effectors such as phosphodiesterases (*Pde10a* and *Pde4d*), protein kinase G (*Prkg2*), protein kinase C (*Prkca*), and the further downstream calcineurin genes (*Ppp3r1* and *Ppp3ca*). These suggest their involvement in GPCR signaling, which regulates cell invasion, mesenchymal transition, formation of polarity, etc. In these clusters, we also find cytoskeleton and cell adhesion associated proteins (*Actr2, Actr3, Arhgef2, Arhgap17*, etc.), and membrane receptor downstream effectors (*Rac1, Map2k2*, etc.) These genes are relevant to the function of cell migration, adhesion, and survival.

Next, we examined the genes with adaptively up-regulated expression in HA-S (clusters 7, 8, and 9). As expected from KEGG analysis, these clusters are overwhelmingly composed of ribosomal-related genes. We also find mitochondrial-related genes in these clusters, including *Atp5g2* and *Sem1*. Since these genes are important for cellular growth, we also examined other adaptive genes in HA-S and found that cellular growth genes, including nucleotide metabolism gene *Tk1*, RNA polimerase subunits (*Polr2j, Polr1d*) and cell cycle gene (*Pcna*) have adaptively up-regulated expression in these sublines.

The functions of genes with adaptively up-regulated expression suggested that HA-R and HA-S sublines are related to the classic “invasive” and “proliferative” phenotypes of melanoma, respectively ^29^. It has been shown that these two phenotypes of melanoma are controlled by two distinct Wnt^6^ pathways. The invasive phenotype is controlled by non-canonical Wnt signaling, which activates PKC/Ca^2+^ signaling to promote cell migration and invasion. The proliferative phenotype is controlled by the canonical Wnt pathway, which is mediated through *β*-catenin to activate Mitf expression, resulting in the induction of genes for cell growth and proliferation ^30,31^. Therefore, we examined if the genes with adaptive expression are involved in the Wnt signaling pathways in a subline-group dependent way. As summarized in Figure 5, invasion-associated genes in clusters 0, 1, and 2 (e.g. G protein subunits, PLC, PKC genes, PKG, calcineurin genes, and actin remodeling genes) are involved in the non-canonical Wnt signaling. In contrast, proliferation-associated genes in clusters 7 to 10 (e.g. ribosomal and mitochondrial genes; cell cycle and RNA polymerase genes) are more involved in the canonical Wnt signaling.

**Figure 5:**
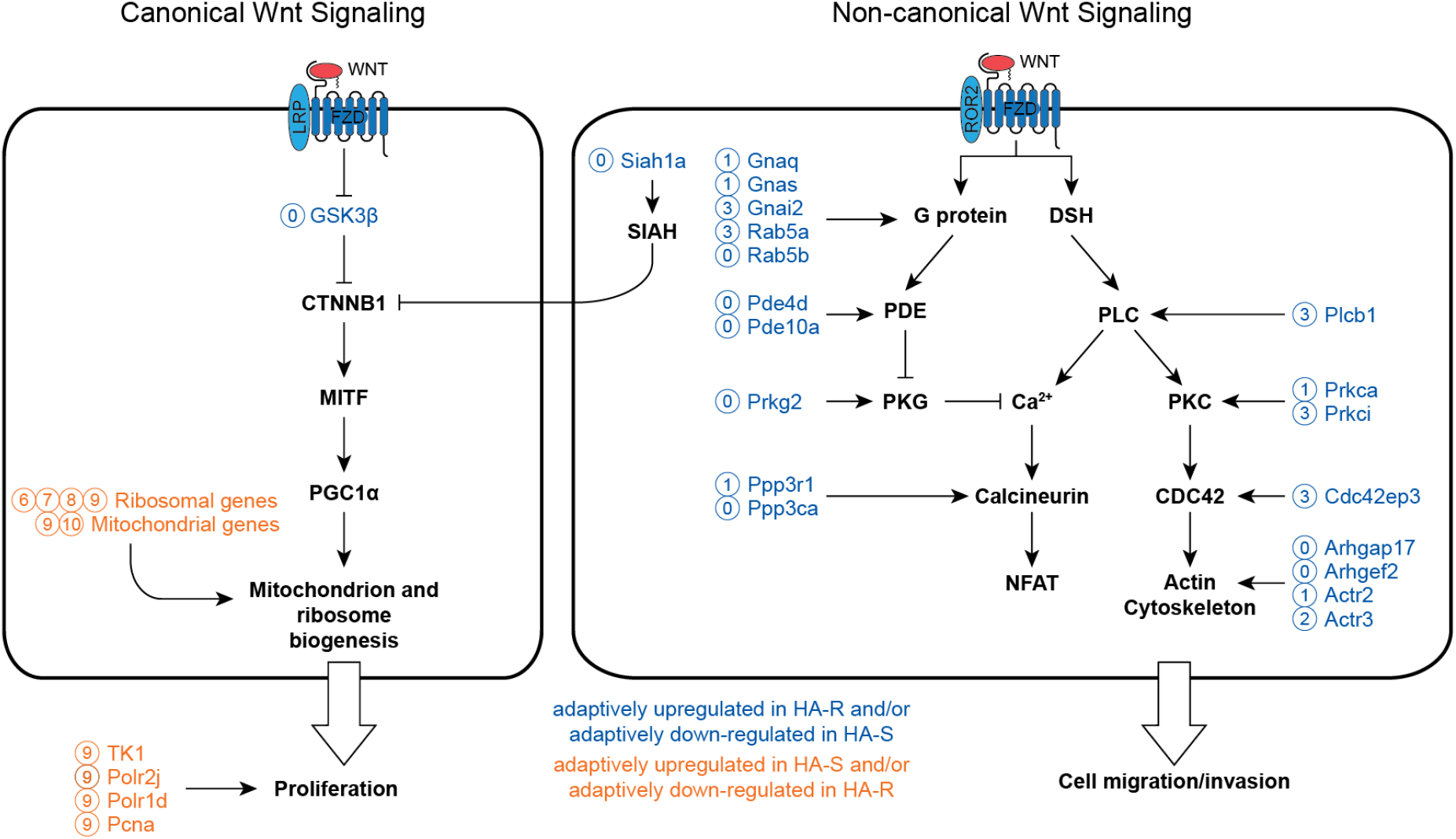
WNT signaling in melanoma. The effectors and signal transduction in each pathway are shown in bold black font. The signaling effector genes that exhibit adaptive expression in high aggression and resistant (HA-R) and high aggression and sensitive (HA-S) are indicated in blue and orange text, respectively. The genes are labeled with the cluster they belong to in the heatmap of Figure 4b. **Left** Canonical WNT signaling. The WNT ligand (e.g. *WNT1, WNT3A*) binds to the composite receptor, *LRP* and Frizzled (*FZD*), resulting in the inhibition of *GSK3β* and activation of *β*-catenine (*CTNNB1*). *CTNNB1* activates *MITF*, which induces the expression of *PGC1α*, ribosomal genes, and melanocytic differentiation genes such as *DCT*, tyrosinase (*TYR*), *TRP1*, etc. *PGC1α* drives the biogenesis of mitochondria. The overall effect is to promote cell differentiation, growth and proliferation. **Right** Non-canonical WNT signaling. The WNT ligand (e.g. *WNT5A, WNT11B*) binds to the composite receptor, *ROR* and *Fzd*, resulting in the activation of calcium signaling through G proteins, phosphodiesterase (PDE), and protein kinase G (PKG), or through dishevelled (DSH) and phospholipase C (PLC). The released calcium (Ca^2+^ will bind to and activate calcineurin, resulting in the NFTA. The PLC signaling may also activate protein kinase C (PKC), which in turn promote cell migration and invasion though CDC42 and cytoskeleton reorganizing genes. The overall effect is to promote epithelial-mesenchymal transition (EMT), cell migration and invasion.

We also found histone modification enzyme genes with adaptively up-regulated expression in HA-R and adaptively down-regulated expression/no-selection in HA-S, including *Kdm1a, Ezh1* (cluster 0), *Kdm5a, Smyd4* (cluster 1), and *Kdm4b, Smyd2* (cluster 2), and *Smyd3* (cluster 10). Since histone modification can determine the activation or inactivation of transcription ^32^, these genes may serve as a “switch” of specific gene modules and therefore have expression undergoing selection. In fact, Siah and Kdm genes have been shown to switch cells from canonical to non-canonical WNT signaling^33^.

In summary, the genes with adaptively up-regulated expression in the high aggression and resistant sublines are associated with cell invasion and non-canonical Wnt signalling. In contrast, the genes with adaptively up-regulated expression in the high aggression and sensitive sublines are associated with cell proliferation and canonical Wnt signaling. Taken together, these results imply that gene modules controlling specific functions and transcriptional states of the cells are subject to selection during melanoma subclonal evolution.

### 2.4 Genes with adaptively evolving expression underline therapeutic responses

In the above analysis, we identified genes with adaptive expression in three different groups of sublines that represent subclones displaying diverse phenotypes. Presumably, the expression of these genes was selected for by the tumor microenvironment, such as immune response and competition of nutrients. These selective pressures gave rise to the distinct phenotypes of the sublines, particularly in vivo growth rate and response to immunity, in a process of adaptation.

However, the 24 sublines isolated in these experiments analysed are expected to be only a sample of all subclones present in the parental line B2905. The emergence of distinct evolutionary related groups (high aggression and resistant (HA-R), high aggression and sensitive (HA-S), low aggression and sensitive (LA-S)) in a randomly selected set of sublines suggests that these groups are representative of cancer evolution. In order to validate if the genes identified to have adaptive expression based on our evolutionary analysis of these sublines represent a general pattern in tumors evolved from the parental line, we examined the relationship between the expression of these genes and the response to ICB treatment of tumors derived from the parental B2905 cell line. If the adaptive expression of the identified genes generalizes across subclones, we should see that the expression of those genes corresponds to the response of the tumor to treatment.

The response of tumors derived from the parental B2905 cell line to ICB treatment was determined in a preclinical study described previously ^34^. Briefly, mice bearing tumors grown from implanted parental B2905 cells were subjected to anti-CTLA-4 or control IgG treatment. After the completion of the dosing, tumors that continuously increased or decreased/remained stable in size for a pre-defined period of time were identified as non-responder and responder tumors, respectively (Figure 6b) and harvested for bulk RNA sequencing. We use a total of 25 mice (14 responders, 6 non-responders, and 5 control).

**Figure 6:**
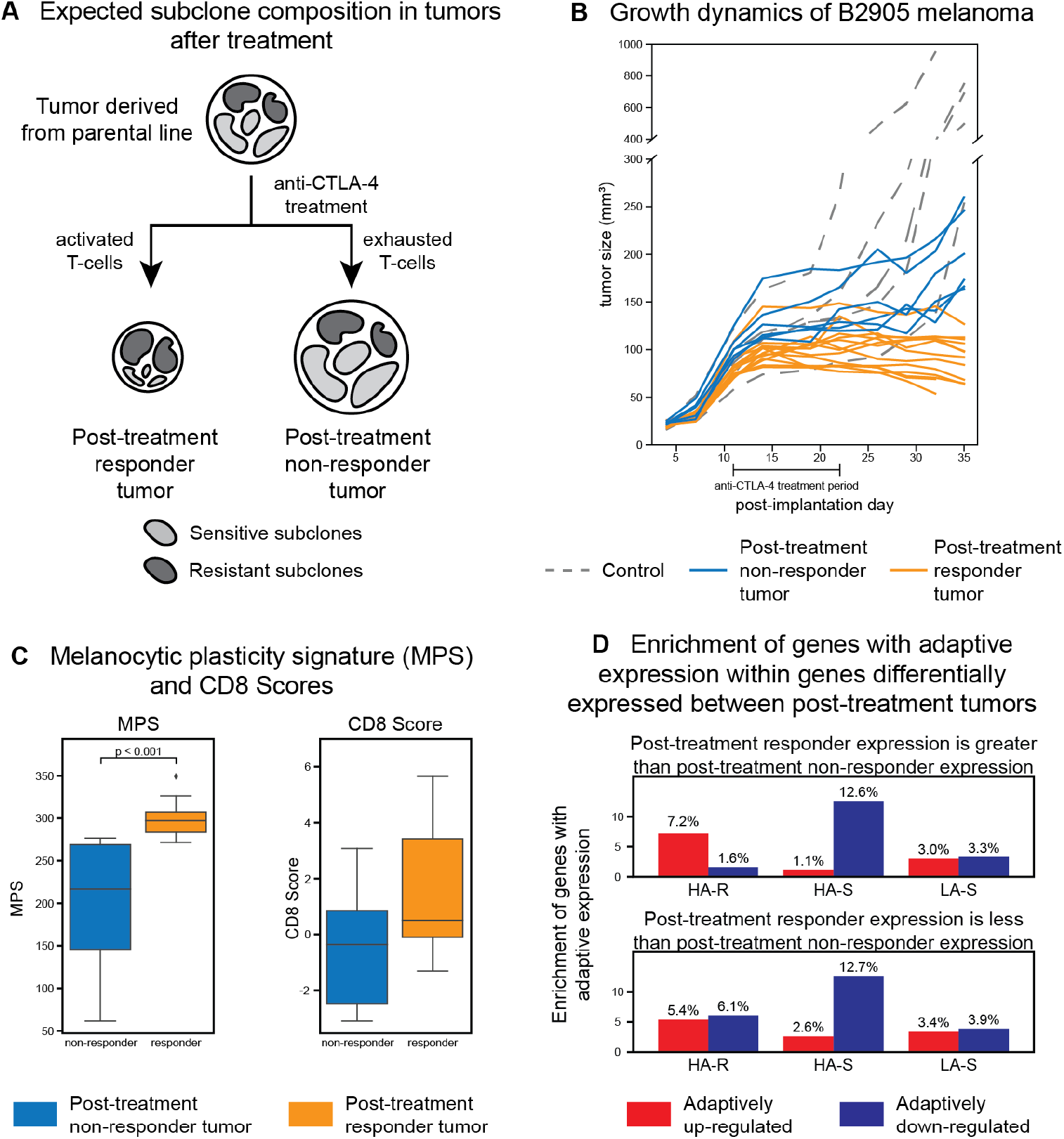
**(a)** The hypothesis of subclonal responses to immunotherapy in melanoma. Parental tumors that contain all subclones are treated with anti-CTLA-4. The therapy could induce effective immunity, resulting in tumor response by reducing the size of the sensitive subclones (light gray) and selecting resistant subclones (dark gray), or fail to induce effective immunity, which results in no response and growth of all subclones. **(b)** Growth of 5 tumors under IgG2b (control; dashed gray lines) and 20 tumors under anti-CTLA-4 treatment (orange and blue lines). The *x*-axis is the day after implantation and the *y*-axis is the size of the tumor in mm^3^. Four doses were given in the period of day 11 to 22 (bar under *x*-axis). The anti-CTLA-4-treated tumors can be divided into 14 responders, where the tumor stopped growing (orange), and 6 non-responders, where the tumor continued to grow (blue). **(c)** The MPS (left) and CD8 (right) scores of the post-treatment responder and non-responder tumors (see text). **(d)** The enrichment of genes with adaptive expression in the genes that are differentially expressed between post-treatment responder and non-responder tumors. The top plot shows genes with higher expression in the post-treatment responder tumors and the bottom plot shows genes with higher expression in the non-responder tumors. The *y*-axes shows the percentage of the differentially expressed genes that have adaptive expression. The *x*-axis shows the different groups of sublines. Each subline group is separated into genes with adaptively up-regulated (red, left) and adaptively down-regulated expression (blue, right).

Tumors that effectively respond to ICB treatment are expected to exhibit higher levels of therapy-induced immunity, reducing the cellular population to resistant subclones after treatment (Figure 6a). To test if the post-treatment responder tumors are indeed dominated by resistant subclones, we evaluated the melanocytic plasticity signature (MPS) score and CD8 expression of the treated tumors (Figure 6c). A high MPS score indicates higher undifferentiated state that is associated with resistance to immunotherapy^23^ and has been used in previous studies to examine treatment resistance^15^. The CD8 score (average of CD8a and CD8b gene expression) has been used to quantify infiltration of effector T cells into tumors, indicating the degree of inflammatory responses^35^. As compared to the post-treatment non-responder tumors, the responder tumors have significantly higher MPS scores (independent t-test p-value *<* 0.001) as well as higher (though not significant) CD8 scores, suggesting therapy-induced effective inflammatory response. This confirms that effective immune response reduces the responder tumors to more undifferentiated, resistant subclones. The resulting resistant subclones are likely to include a broader set of subclones than just those isolated for the single cell analysis. We can test whether genes identified as having adaptive expression using the representative set of sublines are related to immunotherapy response in this wider context. If the identified adaptivity is characteristic of immunotherapy response, then the post-treatment responder tumors should be enriched in genes with adaptive expression in the HA-R sublines because of the dominance of resistant subclones in those tumors. Specifically, when a gene has high expression in the post-treatment responder tumors, that gene is expected to have adaptively up-regulated expression in HA-R sublines, and when a gene has low expression in the responder tumors, that gene is expected to have adaptively down-regulated expression in HA-R sublines.

To investigate the enrichment of genes with adaptive expression in the clonal sublines, we first performed differentially expressed gene (DEG) analysis between post-treatment responder and non-responder tumors using DESeq2^36^. Of the 2,268 DEGs that are also in the single cell dataset used to test for adaptivity, 642 and 1,626 have higher relative mean expression in the post-treatment responder and non-responder tumors, respectively.

We then examined the enrichment of genes with adaptive expression within the DEGs by calculating the percentage of adaptively expressed genes in the DEGs (Figure 6d). As expected, the DEGs that are more highly expressed in the post-treatment responder tumors are preferentially enriched with genes with adaptively up-regulated expression in the HA-R sublines relative to those with adaptively down-regulated expression (7.2% versus 1.6%). In contrast, DEGs more highly expressed in post-treatment non-responder tumors are preferentially enriched with genes with adaptively down-regulated expression in the HA-R sublines relative to those with adaptively up-regulated expression (6.1% versus 5.4%), although this trend is less distinct. The differences between these enrichment patterns are statistically significant (chi-squared p-value *<* 0.0001). These patterns demonstrate that the genes identified as having adaptive expression in the HA-R sublines are indeed more highly expressed in the post-treatment responder tumors, which experienced effective treatment and represent a resistant population of cells.

Next, we analyzed the genes with adaptive expression in the HA-S sublines. The DEGs with higher expression in the post-treatment responder tumors are preferentially enriched with genes with adaptively down-regulated expression relative to those with adaptively upregulated expression (12.6% versus 1.1%). The genes with higher expression in the post-treatment non-responder tumors also have higher enrichment with genes with adaptively down-regulated expression relative genes with adaptively up-regulated expression (12.7% versus 2.6%). This trend may be because of the likely heterogeneity of the post-treatment non-responder tumors (Figure 6a). These enrichment differences are statistically significant (chi-squared p-value *<* 0.05). Genes with adaptive expression in the LA-S sublines do not show significant preferential enrichment. These analyses further support the view that genes identified to have adaptively up-regulated expression in HA-R are broadly related to the invasive and ICB resistant tumor phenotype.

## 3 Discussion

### Summary

In this study we utilized Brownian motion and Ornstein-Uhlenbeck processes to model adaptive evolution of gene expression in mouse melanoma tumors. By running our method on clonal sublines derived from a mouse melanoma model exhibiting distinct phenotypic properties, we were able to identify genes with adaptive expression within groups of sublines. Specifically, we focused on sublines that varied in their growth rate and response to anti-CTLA-4 immunotherapy treatment when implanted individually into distinct but genetically identical mice.

We found that about 7-15% of the genes have expression that can be considered to be adaptive within the different subline groups. The genes with adaptive expression in sublines that had high aggression and were resistant to ICB therapy (HA-R) are associated with cell invasion, cell migration, adhesion, and survival. In contrast, the genes with adaptive expression in the sublines that had high aggression and were sensitive to ICB therapy (HAS) are associated with ribosomal and mitochondrial processes, indicating cell growth. These two groups of genes represent the known binary invasive and proliferative phenotypes of melanoma. Additionally, the genes with adaptively up-regulated expression in the HA-R sublines are associated with non-canonical Wnt signaling, while the genes with adaptively up-regulated expression in the HA-S sublines are associated with canonical Wnt signaling, further supporting the link with the binary phenotypes in melanoma ^29^.

We also evaluated the genes with adaptive expression in tumors derived from the parental line and treated with anti-CTLA-4 immunotherapy. We found that tumors with therapy-induced immunity were enriched with genes with adaptive expression in HA-R. This is consistent with our previous study showing that resistance of melanoma to ICB is associated with the precursor states ^23^. Our results suggest that genes in functional modules are subjected to selection together. Such gene modules are usually involved in the developmental process^37^. For example, we identify selection on gene expression related to non-canonical and canonical Wnt signaling genes. These genes are involved in the migration and maturation phase of melanocyte precursors ^38^. Our findings also indicate that selection of gene modules could be achieved by the adaptively up-regulated expression of histone modification enzyme genes, which can serve as genetic switches of gene groups.

### Stochastic models of gene expression expand on traditional mutation and sequence based methods

In contrast to previous tumor evolution analyses that focus on mutations, our analysis focuses on the evolution of gene expression. The insights from expression-based evolutionary analysis using a mutation-based phylogeny are complementary to analyses using mutations alone. In particular, we found that mutated genes are not more likely to have adaptive expression, suggesting that our results would not be identifiable if solely using mutations. This may imply the distinct roles of genetic and non-genetic drivers of subclonal evolution. In addition, we performed analysis with traditional dN/dS methods, which are used to determine adaptivity in context of sequence evolution. Using counting-based methods and dNdScv ^39^, we found a potential small positive selection across the entire genome, but the sparsity of mutational data excluded detailed gene-wise analysis. Thus, the modeling of gene expression evolution in cancer provides unique insights into tumor evolution that an analysis based on mutations alone would miss. See the Supplementary Sections 3 and 4 for the details of these analyses.

### Limitations of the study and opportunities for future work

There are limitations of our study that open possibilities for future work. First, because of the noise in single-cell data, our method, like differential expression and sequence-based dN/dS methods, is unable to distinguish weak signals of constrained evolution from neutral evolution, as shown in our simulation studies. This makes it unsurprising that we only identify a few genes with constrained expression evolution in the biological data (these results are further discussed in the Supplementary Section 2). However, we cannot rule out the possibility that tumors evolve with limited constrained evolution. This question could be further investigated by leveraging bulk RNA sequencing, which has less noise, in conjunction with single-cell data.

Second, when analyzing sublines from one regime (e.g., HA-R), the sublines in the other regimes (e.g., HA-S and LA-S) are considered as part of the background. Our results then must be considered comparatively. This could result in genes that are only in fact adaptively up-regulated in one regime being also labeled as adaptively down-regulated in the others. To help avoid comparative results, Ornstein-Uhlenbeck (OU) models allow for more than two optima to be estimated. In order to address whether using only two optima impacted our results, we ran experiments with such further divisions, considering the case where the HA-R, the HA-S, and the rest of the sublines all have separate optima. The results of this three regime experiment were similar to the two-regime results, giving us increased confidence that our results using two optima can be generalized. Using more optima increases the number of parameters to estimate and reduces the power of the stochastic methods, resulting in fewer genes being properly estimated. Designing new methods that can better optimize over additional optima would allow us to further support our results and enable future studies examining multiple groups of sublines.

Further, previous studies in species evolution have suggested that the size and branch length of the evolutionary tree can impact results of Brownian motion and Ornstien-Uhlenbeck methods, where increasing the number of leaves and lengths of branches improved model performance ^22,40,17^. While our tree has 23 leaves, more than other studies of species evolution ^26,17,21^, it is possible that adding more sublines could provide even further insights.

Despite these limitations, our method was able to identify genes with adaptive expression, and our subsequent analysis reveals that these adaptive genes were associated with specific pathways directly linked to the subline phenotype. We were unable to identify these genes from dN/dS analyses or mutations alone, demonstrating the benefits of gene expression evolution analyses for studying subclonal evolution and response to treatment. Applying stochastic methods to other cancer types can reveal further understanding of general and cancer type-specific adaptation.

### 3.1 Conclusions

In conclusion, our study demonstrates the efficacy of using stochastic processes for modeling gene expression evolution of tumor sublines in context of uncovering clone-specific adaptation. Applying our methodology to mouse melanoma sublines revealed adaptive evolution patterns consistent with the subline phenotype and response to treatment. Specifically, sublines that were resistant or sensitive to treatment had phenotype-specific profiles of genes with adaptive expression. This finding led us to discover pathways undergoing selection, specifically Wnt signaling, that vary between the sublines with contrasting phenotypes. Subsequent gene expression analysis of tumors grown from the parental line and subjected to anti-CTLA-4 treatment showed that the genes identified to have adaptive expression based on our analysis of isolated sublines are broadly associated with the corresponding phenotype. Our methodology for analyzing clonal sublines identified phenotype-switching signaling that could not be revealed by “snapshot” analysis, such as differential expression or dN/dS analyses.

Our results demonstrate that analysis of single-cell sequencing data using stochastic models can identify genes and functional modules with expression undergoing adaptation, which is key information in tumor evolution and heterogeneity. Genes with adaptive expression can inform candidate gene selection for targeted therapies for tumors exhibiting specific phenotypic properties.

## Author contributions

M.G.H., E.K.M., C.-P.D, and T.M.P. were involved in conceptualization of the project. M.G.H., C.-P.D, S.M., F.R.M., and E.P.G. performed data curation. M.G.H., E.K.M., C.- P.D., and T.M.P. performed the investigation. C.G., A.S., and E.P.G. created the in vitro and in vivo models. M.G.H., S.P., C.-P.D., and T.M.P. developed the methodology of the project. M.G.H., C.-P.D., and T.M.P. performed the formal analysis. M.G.H. and S.P. developed the software used in this project. M.G.H., S.P., and C.-P.D. performed validation of the computational models. M.G.H. and C.-P.D. created visualizations. G.M. provided the melanoma cell line B2905 and its 24 sublines. M.G.H., E.K.M., C.-P.D., and T.M.P. wrote the original draft of the manuscript. S.P., C.S., and G.M. reviewed and edited the manuscript. E.K.M. and T.M.P. supervised the computational analysis; C.-P.D. supervised the preclinical study. T.M.P. and G.M. acquired funding for the project.

## Acknowledgments

M.G.H. and T.M.P are supported by the NLM Intramural Research Program. S.P. is supported by the NEI Intramural Research Program. F.R.M., S.M., C.G., E.P.G., C.S., G.M., and C.-P.D. are supported by the NCI Intramural Research Program. The research of E.P.G., G.M., and C.-P.D. is supported in part by FLEX Synergy Award from the NCI Center for Cancer Research. E.P.G is supported by Spanish Agencia Estatal de Investigación and Ministerio de Ciencia e Innovacion (PID2022-141113OA-I00) and a MRA Young Investigator Award (#1037420) by Melanoma Research Alliance. E.K.M. is supported by the State of Maryland. Computational experiments were performed on the compute cluster for the Center for Bioinformatics and Computational Biology at the University of Maryland, College Park. The authors thank Ms. Sung Chin and Cari Smith (Laboratory Animal Science Program, Frederick National Laboratory for Cancer Research) for performing mouse studies, and Dr. Yongmei Zhao (CCR Sequencing Facility Bioinformatics Group, Frederick National Laboratory for Cancer Research) for assistance coordinating the bulk treatment data.

## STAR Methods

### Lead contact

Further information and requests for resources and reagents should be directed to and will be fulfilled by the lead contact, Teresa M. Przytycka.

### Materials availability

No newly generated material is associated with this paper.

### Data and code availability

#### Data

Single-cell RNA-seq data have been deposited at GEO and are publicly available as of the date of publication. Accession number is GSE255484.

#### Code

All original code has been deposited at GitHub and is publicly available as of the date of publication. https://github.com/ncbi/nongenetic-evolution-of-tumor-subclones

#### Additional information

Any additional information required to reanalyze the data reported in this paper is available from the lead contact upon request.

### Experimental Model and Study Participant Details

For M4 melanoma model, pups of hepatocyte growth factor (HGF)-transgenic C57BL/6 mice received UV irradiation at postnatal day 3, and melanoma would emerged in 6-8 months. A melanoma was harvested from a male mouse to make B2905 cell line ^41^. B2905 melanoma carries spontaneous hotspot mutations in *Kras, Gnaq*, and *Sf3b1* ^23^. The M4 melanoma model for preclinical study was built by implanting B2905 cells into C57BL/6 mice subcutaneously.

The development of the 24 clonal sublines was described previously ^24^. In brief, 24 single cells were randomly selected and isolated from the parental B2905 mouse melanoma line, which was generated ^23^ and cultured in our laboratory, by cell sorting without any labeling. They were expanded in culture to become individual clonal sublines, C1 to C24. Their analyses of whole exome, whole transcriptomics, and single-cell full transcripts were also described by^24^.

For the preclinical study of B2905 melanoma response to anti-CTLA-4 (Section 2.4), Wildtype female C57BL/6 at the age of 6-8 weeks old were purchased from The Charles River at NCI Frederick. Mice at Laboratory Animal Science Program, National Frederick Laboratory for Cancer Research were housed in micro-isolator and individually ventilated cages supplied with acidified water and fed 5053 Irradiated Picolab Rodent Diet 20 lab diet. Temperature for laboratory mice in our facility is maintained between 18–23°C with 40–60% humidity. All animal procedures are conducted in laminar flow hoods. All personnel are required to wear scrubs and lab coat, mask, hair net, shoe covers, and disposable gloves upon entering the animal rooms. A 12 light/12 dark cycle was used for the mice. The animal study protocol (16-007) was approved by the Animal Care and Use Committee at NCI Frederick.

In all, 1.0× 10^6^ B2905 cells were implanted into the flank of 40 female C57Bl/6 mice, aged 6-8 weeks. The tumor size was measured twice a week. When tumors reached 75 mm^3^ of size by average, mice were randomized into three groups to receive treatment of anti-CTLA-4 (Bio X Cell Cat.No. BE0164, clone 9D9; 30 mice), IgG2b isotype control (Bio X Cell Cat.No. BP0086, clone MPC-11; 5 mice), or left untreated (5 mice). The antibody was given to mice at 10 mg/kg i.v., twice a week for two consecutive weeks. After the completion of antibody dosing, tumors with size approximately unchanged or decreasing at the next three measurements were identified as stable disease or responder tumors, respectively, and harvested. For those with size continuously increasing at the next three measurements, they were identified as non-responder and harvested at designated size. In any group, when the tumor reached 2000 mm^3^ or ulcerated, or mice showed behavior of sickness, the endpoint was achieved, the mice would be euthanized by trained personnel with carbon dioxide inhalation in a euthanasia chamber, following the procedure in our animal study protocol. 25 tumors were harvested before reaching the endpoint and subjected to RNA sequencing. The rest were allowed to reach endpoint for observing therapeutic efficacy. After euthanization, the tumor would be harvested for flash freezing in liquid nitrogen or fixation in paraformaldehyde, and the mouse would be examined by necropsy. The flash-frozen tumor would be processed for RNA sequencing.

### Quantification and Statistical Analysis

#### Determining adaptive genes with Brownian motion and Ornstein-Uhlenbeck models

We use three stochastic models to determine adaptivity: a Brownian motion model, an Ornstein-Uhlenbeck model with a single optimum, and an Ornstein-Uhlenbeck model with two optima. These models describe the change of expression for a gene for a subline over time. Brownian motion describes a stochastic process where the gene expression values change by a random value from a normal distribution at each time step *t*:

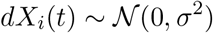

where *X*_*i*_ is the expression value of species *i*, and *σ* is the standard deviation of the normal distribution.

The Ornstein-Uhlenbeck model is a generalization of Brownian motion that includes a pull to an “optimum” value at each time step:

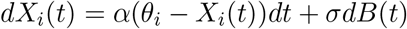

where *θ*_*i*_ is the optimum value for species *i, α* is the influence of the optimum on the change, and *dB*(*t*) is change from Brownian motion. When *α* = 0, this model simplifies to Brownian motion.

We extend this to include multiple replicate values for each gene. To model this, we add a noise term to the species value:

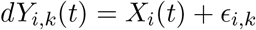

where *Y*_*i,k*_ is the replicate value for species *i*, replicate *k, ϵ*_*i,k*_ ∼ N (0, *γσ*^2^) is a Gaussian variable with mean 0 and variance γ σ^2^, which represents the within-species variability.

Parameters *σ, α, θ*_*i*_, and *γ* are estimated using the maximum likelihood procedure described in the original EvoGeneX study ^17^. Initial values of the parameters *α* and *γ* were set to 0.1 and 0.01, respectively, as they are shown to work well in the original study ^17^.

### Function and pathway enrichment analysis

We use clusterProfiler ^42^ in R to determine enrichment of KEGG terms in genes using over-representation analysis. In the single cell analysis, we determine enrichment of the genes with adaptive expression for each of the regimes. The background gene set for this analysis is all the genes in the single cell data. In the bulk validation analysis, we use the genes that appear both in the single cell data and the bulk data for the background set. We correct p-values with using Benjamini-Hochberg false discovery rate correction and consider pathways with a corrected p-value *<* 0.05 as significant.

### Clustering of genes with adaptive expression

We perform k-means clustering on the genes with adaptive expression in at least one of the three regimes we analyzed. In order to do this, we assign each gene a value for each regime based on the BH-FDR corrected p-value for that regime, resulting in a set of three values for each gene. Specifically, for each regime *r*, a gene is assigned the following value:

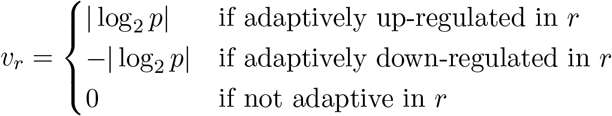

The genes are clustered based on these triplet values (ie, the vector [*v*_HA-R_, *v*_HA-S_, *v*_LA-S_]. We ran k-means clustering with 2 to 20 clusters and chose to use 12 clusters because it had the lowest silhouette score.

### Differential expression analysis

We used DESeq2^36^ to perform differential expression analysis between post-treatment responder and non-responder tumors. We used as input the count matrix of the gene expression and ran DESeq2 with the default parameters. P-values were corrected for multiple hypothesis testing using the Benjamini-Hochberg false discovery rate method and genes with corrected p-values *<* 0.05 were considered significant.

## Footnotes

1 The M4 mouse melanoma model represents RAS-mutated human melanoma with a transitory differentiation subtype. See Methods for detailed description.

2 Each clonal subline was grown from a single cell isolated from the B2905 cell line ^24^.

3 Subline C2 was excluded after data filtering ^15^.

4 While partitioning the tree to have more than two optima is possible, the addition of optima values to estimate reduces the power of the method, so here we only consider two optima.

5 HA-S was also enriched in the new Coronavirus pathway (hsa05171), but we found that this is caused by the enrichment in ribosomal genes ^28^ that constitute 89 out of 232 genes in this pathway so we do not include this pathway in the discussion.

6 In accordance with organism-specific formatting guidelines (https://www.ncbi.nlm.nih.gov/genbank/eukaryotic_genome_submission_annotation/), the mouse gene symbol begins with an uppercase letter, followed by lowercase letters, and human gene symbol uses all uppercase letters.

## References

[1] Nowell, P.C. (1976). The clonal evolution of tumor cell populations: acquired genetic lability permits stepwise selection of variant sublines and underlies tumor progression. Science, 194(4260), 23–28. 10.1126/science.959840.

[2] Vendramin, R., Litchfield, K., and Swanton, C. (2021). Cancer evolution: Darwin and beyond. EMBO J, 40(18), e108389. 10.15252/embj.2021108389.

[3] Scott, J. and Marusyk, A. (2017). Somatic clonal evolution: a selection-centric perspective. Biochim Biophys Acta Rev Cancer, 1867(2), 139–150. 10.1016/j.bbcan.2017.01.006.

[4] Beerenwinkel, N., Schwarz, R.F., Gerstung, M., and Markowetz, F. (2015). Cancer evolution: mathematical models and computational inference. Syst Biol, 64(1), e1–e25. 10.1093/sysbio/syu081.

[5] Williams, M.J., Sottoriva, A., and Graham, T.A. (2019). Measuring clonal evolution in cancer with genomics. Annu Rev Genomics Hum Genet, 20, 309–329. 10.1146/annurev-genom-083117-021712.

[6] Caiado, F., Silva-Santos, B., and Norell, H. (2016). Intra-tumour heterogeneity–going beyond genetics. FEBS J, 283(12), 2245–2258. 10.1111/febs.13705.

[7] Yang, D., Jones, M.G., Naranjo, S., Rideout, W.M., Min, K.H.J., Ho, R., Wu, W., Replogle, J.M., Page, J.L., Quinn, J.J., et al. (2022). Lineage tracing reveals the phylodynamics, plasticity, and paths of tumor evolution. Cell, 185(11), 1905–1923.e25. 10.1016/j.cell.2022.04.015.

[8] Liu, Y., Li, X.C., Mehrabadi, F.R., Schäffer, A.A., Pratt, D., Crawford, D.R., Malikić, S., Molloy, E.K., Gopalan, V., Mount, S.M., et al. (2023). Single-cell methylation sequencing data reveal succinct metastatic migration histories and tumor progression models. Genome Res, 33(7), 1089–1100. 10.1101/gr.277608.122.

[9] Saelens, W., Cannoodt, R., Todorov, H., and Saeys, Y. (2019). A comparison of single-cell trajectory inference methods. Nat Biotechnol, 37(5), 547–554. 10.1038/s41587-019-0071-9.

[10] La Manno, G., Soldatov, R., Zeisel, A., Braun, E., Hochgerner, H., Petukhov, V., Lidschreiber, K., Kastriti, M.E., Lönnerberg, P., Furlan, A., et al. (2018). Rna velocity of single cells. Nature, 560(7719), 494–498. 10.1038/s41586-018-0414-6.

[11] Matsumoto, H. and Kiryu, H. (2016). Scoup: a probabilistic model based on the ornstein-uhlenbeck process to analyze single-cell expression data during differentiation. BMC bioinformatics, 17(1), 1–16. 10.1186/s12859-016-1109-3.

[12] Wang, L., Fan, J., Francis, J.M., Georghiou, G., Hergert, S., Li, S., Gambe, R., Zhou, C.W., Yang, C., Xiao, S., et al. (2017). Integrated single-cell genetic and transcriptional analysis suggests novel drivers of chronic lymphocytic leukemia. Genome Res, 27(8), 1300–1311. 10.1101/gr.217331.116.

[13] McCarthy, D.J., Rostom, R., Huang, Y., Kunz, D.J., Danecek, P., Bonder, M.J., Hagai, T., Lyu, R., Wang, W., Gaffney, D.J., et al. (2020). Cardelino: computational integration of somatic clonal substructure and single-cell transcriptomes. Nat Methods, 17(4), 414–421. 10.1038/s41592-020-0766-3.

[14] Shafighi, S.D., Kie lbasa, S.M., Sepúlveda-Yáñez, J., Monajemi, R., Cats, D., Mei, H., Menafra, R., Kloet, S., Veelken, H., van Bergen, C.A., et al. (2021). Cactus: integrating clonal architecture with genomic clustering and transcriptome profiling of single tumor cells. Genome Med, 13, 1–16. 10.1186/s13073-021-00842-w.

[15] Mehrabadi, F.R., Marie, K.L., Pérez-Guijarro, E., Malikić, S., Azer, E.S., Yang, H.H., Kızılkale, C., Gruen, C., Robinson, W., Liu, H., et al. (2021). Profiles of expressed mutations in single cells reveal subclonal expansion patterns and therapeutic impact of intratumor heterogeneity. bioRxiv. 10.1101/2021.03.26.437185.

[16] Bastide, P., Soneson, C., Stern, D.B., Lespinet, O., and Gallopin, M. (2023). A phylogenetic framework to simulate synthetic interspecies rna-seq data. Mol Biol Evol, 40(1). 10.1093/molbev/msac269z.

[17] Pal, S., Oliver, B., and Przytycka, T.M. (2023). Stochastic modeling of gene expression evolution uncovers tissue-and sex-specific properties of expression evolution in the drosophila genus. J Comput Biol, 30(1), 21–40. 10.1089/cmb.2022.0121.

[18] Bedford, T. and Hartl, D.L. (2009). Optimization of gene expression by natural selection. Proc Natl Acad Sci USA, 106(4), 1133–1138. 10.1073/pnas.0812009106.

[19] Bertram, J., Fulton, B., Tourigny, J.P., Peña-Garcia, Y., Moyle, L.C., and Hahn, M.W. (2023). Cagee: computational analysis of gene expression evolution. Mol Biol Evol, 40(5), msad106. 10.1093/molbev/msad106.

[20] Butler, M.A. and King, A.A. (2004). Phylogenetic comparative analysis: a modeling approach for adaptive evolution. Am Nat, 164(6), 683–695. 10.1086/426002.

[21] Brawand, D., Soumillon, M., Necsulea, A., Julien, P., Csárdi, G., Harrigan, P., Weier, M., Liechti, A., Aximu-Petri, A., Kircher, M., et al. (2011). The evolution of gene expression levels in mammalian organs. Nature, 478(7369), 343–348. 10.1038/nature10532.

[22] Rohlfs, R.V., Harrigan, P., and Nielsen, R. (2014). Modeling gene expression evolution with an extended ornstein-uhlenbeck process accounting for within-species variation. Mol Biol Evol, 31(1), 201–211. 10.1093/molbev/mst190.

[23] Pérez-Guijarro, E., Yang, H.H., Araya, R.E., El Meskini, R., Michael, H.T., Vodnala, S.K., Marie, K.L., Smith, C., Chin, S., Lam, K.C., et al. (2020). Multimodel preclinical platform predicts clinical response of melanoma to immunotherapy. Nat Med, 26(5), 781–791. 10.1038/s41591-020-0818-3.

[24] Gruen, C., Yang, H.H., Sassano, A., Wu, E., Gopalan, V., Marie, K.L., Castro, A., Mehrabadi, F.R., Wu, C.H., Church, I., et al. (2023). Melanoma clonal subline analysis uncovers heterogeneity-driven immunotherapy resistance mechanisms. bioRxiv, pp. 2023– 04. 10.1101/2023.04.03.535074.

[25] Hansen, T.F. (1997). Stabilizing selection and the comparative analysis of adaptation. Evolution, 51(5), 1341–1351. 10.1111/j.1558-5646.1997.tb01457.x.

[26] Chen, J., Swofford, R., Johnson, J., Cummings, B.B., Rogel, N., Lindblad-Toh, K., Haerty, W., Di Palma, F., and Regev, A. (2019). A quantitative framework for characterizing the evolutionary history of mammalian gene expression. Genome Res, 29(1), 53–63. 10.1101/gr.237636.118.

[27] Price, P.D., Palmer Droguett, D.H., Taylor, J.A., Kim, D.W., Place, E.S., Rogers, T.F., Mank, J.E., Cooney, C.R., and Wright, A.E. (2022). Detecting signatures of selection on gene expression. Nat Ecol Evol, pp. 1–11. 10.1038/s41559-022-01761-8.

[28] Jiao, L., Liu, Y., Yu, X.Y., Pan, X., Zhang, Y., Tu, J., Song, Y.H., and Li, Y. (2023). Ribosome biogenesis in disease: new players and therapeutic targets. Sig Transduct Target Ther, 8(1), 15. 10.1038/s41392-022-01285-4.

[29] Rambow, F., Marine, J.C., and Goding, C.R. (2019). Melanoma plasticity and phenotypic diversity: therapeutic barriers and opportunities. Genes Dev, 33(19-20), 1295–1318. 10.1101/gad.329771.119.

[30] Chauhan, J.S., Hölzel, M., Lambert, J.P., Buffa, F.M., and Goding, C.R. (2022). The mitf regulatory network in melanoma. Pigment Cell Melanoma Res, 35(5), 517–533. 10.1111/pcmr.13053.

[31] Webster, M.R., Kugel III, C.H., and Weeraratna, A.T. (2015). The wnts of change: How wnts regulate phenotype switching in melanoma. Biochim Biophys Acta Rev Cancer, 1856(2), 244–251. 10.1016/j.bbcan.2015.10.002.

[32] Ninova, M., Fejes Tóth, K., and Aravin, A.A. (2019). The control of gene expression and cell identity by h3k9 trimethylation. Development, 146(19), dev181180. 10.1242/dev.181180.

[33] Wend, P., Holland, J.D., Ziebold, U., and Birchmeier, W. (2010). Wnt signaling in stem and cancer stem cells. In Semin Cell Dev Biol (Elsevier), volume 21, pp. 855–863. 10.1016/j.semcdb.2010.09.004.

[34] Pagadala, M., Sears, T.J., Wu, V.H., Pérez-Guijarro, E., Kim, H., Castro, A., Talwar, J.V., Gonzalez-Colin, C., Cao, S., Schmiedel, B.J., et al. (2023). Germline modifiers of the tumor immune microenvironment implicate drivers of cancer risk and immunotherapy response. Nat Commun, 14(1), 2744. 10.1038/s41467-023-38271-5.

[35] Jiang, P., Gu, S., Pan, D., Fu, J., Sahu, A., Hu, X., Li, Z., Traugh, N., Bu, X., Li, B., et al. (2018). Signatures of t cell dysfunction and exclusion predict cancer immunotherapy response. Nat Med, 24(10), 1550–1558. 10.1038/s41591-018-0136-1.

[36] Love, M.I., Huber, W., and Anders, S. (2014). Moderated estimation of fold change and dispersion for rna-seq data with deseq2. Genome Biol, 15(12), 1–21. 10.1186/s13059-014-0550-8.

[37] Baggiolini, A., Callahan, S.J., Montal, E., Weiss, J.M., Trieu, T., Tagore, M.M., Tischfield, S.E., Walsh, R.M., Suresh, S., Fan, Y., et al. (2021). Developmental chromatin programs determine oncogenic competence in melanoma. Science, 373(6559), eabc1048. 10.1126/science.abc1048.

[38] Gopalan, V., Day, C.P., Pérez-Guijarro, E., Chin, S., Ebersole, J., Smith, C., Simpson, M., Sassano, A., Alves Constantino, M., Wu, E., et al. (2022). Comprehensive single-cell transcriptomic analysis of embryonic melanoblasts uncovers lineage-specific mechanisms of melanoma metastasis and therapy resistance. bioRxiv, pp. 2022–10. 10.1101/2022.10.14.512297.

[39] Martincorena, I., Raine, K.M., Gerstung, M., Dawson, K.J., Haase, K., Van Loo, P., Davies, H., Stratton, M.R., and Campbell, P.J. (2017). Universal patterns of selection in cancer and somatic tissues. Cell, 171(5), 1029–1041. 10.1016/j.cell.2017.09.042.

[40] Cooper, N., Thomas, G.H., Venditti, C., Meade, A., and Freckleton, R.P. (2016). A cautionary note on the use of ornstein uhlenbeck models in macroevolutionary studies. Biol J Linn Soc Lond, 118(1), 64–77. 10.1111/bij.12701.

[41] Noonan, F.P., Zaidi, M.R., Wolnicka-Glubisz, A., Anver, M.R., Bahn, J., Wielgus, A., Cadet, J., Douki, T., Mouret, S., Tucker, M.A., et al. (2012). Melanoma induction by ultraviolet a but not ultraviolet b radiation requires melanin pigment. Nat Commun, 3(1), 884. 10.1038/ncomms1893.

[42] Wu, T., Hu, E., Xu, S., Chen, M., Guo, P., Dai, Z., Feng, T., Zhou, L., Tang, W., Zhan, L., et al. (2021). clusterprofiler 4.0: A universal enrichment tool for interpreting omics data. Innovation (Camb), 2(3), 100141. 10.1016/j.xinn.2021.100141.

