## Supplementary Text for "Stochastic modelling of single-cell gene expression adaptation reveals non-genomic contribution to evolution of tumor subclones"

### Contents

**1 Simulation Studies** **1**

1.1 Data generation . . . . . 1

1.2 Results . . . . . 2

**2 Lack of constrained adaptation** **3**

**3 Mutated genes are not more likely to have adaptive expression** **3**

**4 dN/dS** **4**

**References** **6**

### 1 Simulation Studies

#### 1.1 Data generation

We perform simulation experiments in order to show that EvoGeneX[1] is appropriate for single cell cancer data. We do this by simulating TPM data under neutral, constrained, and adaptive models with high levels of noise that resemble the single-cell RNA sequence data we analyze.

In order to simulate single cell data undergoing different modes of evolution, we first simulate a Brownian motion (BM)/Ornstein-Uhlenbeck (OU) process down the tree with the root value fixed. We use the R library OUwie to model BM and OU processes. Rather than create replicates using a normal distribution as previously used to simulate bulk data[1], we draw from a negative binomial distribution, which is a more standard way to simulate single-cell RNA sequence data[2,3]. The negative binomial has a mean equal to the species value and dispersion parameter $r$, giving the expression value of species $i$, replicate $k$ at time $t$ as

$$Y_{i,k}(t) \sim \mathrm{NB}(X_i(t), r)$$

The variance of the distribution $(\mu + \frac{\mu^2}{r})$ represents the within-species variation, as compared to $\gamma\sigma^2$ when drawing replicates to be from a normal distribution as is expected in EvoGeneX. As a result, rather than having a parameter $\gamma$ that controls the within-species variation, we have a parameter $r$, where as $r$ increases, the within-species variation decreases.

We run simulations over the mutation tree from the 24 subline experiments with the high aggression and resistant (HA-R) regime and use parameter values estimated from the data. We set the expression level at the root to be 150, approximately equal to the mean of the expression values of all the replicates in the single cell dataset. We vary $\sigma^2$ between 0.1%, 1%, and 10% of the expression at the root (0.15, 1.5, and 15) and $\alpha$ between 0.125, 1, and 8, which are similar to the values used in the simulations in Pal et al.[1]. High $\sigma^2$ values represent high between-species variation, and high $\alpha$ values represent a strong pull to the optimum value. We vary $\theta_1/\theta_0$ between 1.02, 1.04, 1.06, 1.08, and 1.1. The $\theta$ values are equidistant from the root value and $\theta_1$ is higher and $\theta_0$ is lower than the root value, respectively. Low $\theta_1/\theta_0$ values represent instances where the optima values are very similar, making the different harder to detect. We vary $r$ between 100, 150, and 200, where 150 reduces the equation to a Poisson distribution, and 100 and 200 represent under and over dispersion, respectively. Low $r$ values represent high within-species variation.

We generate three types of data: 1) neutral data using Brownian motion (900 parameter combinations), 2) constrained data using an OU process with a single optimum (2,700 parameter combinations), and 3) adaptive data using an OU process with two optima using the HA-R regime (13,500 parameter combinations). For each combination of parameters, we simulate 100 genes, each with at most eight replicates with each having a 10% chance of drop-out to resemble the missing data in the real dataset.

Over every dataset, we run EvoGeneX with BM, OU with one optima, and OU with two optima. Running OU processes over the neutral data will demonstrate the false positive rate. Running an OU process with a single optimum over data simulated as constrained and an OU process with two optima over the data simulated as adaptive will demonstrate the true positive rate.

#### 1.2 Results

EvoGeneX correctly predicts approximately 41.7% of the adaptive genes as being adaptive. EvoGeneX incorrectly predicts 4.7% and 3.4% of constrained and neutral genes to be adaptive, respectively. 0.73% of adaptive genes are incorrectly predicted by EvoGeneX as being constrained; no neutral genes are predicted to be constrained. Only 1.63% of the constrained genes are correctly predicted by EvoGeneX as being constrained. We expect this is because the negative binomial overpowers the reduced variation between the species.

46.4% of the adaptive genes, 26.3% of neutral genes, and 0.52% of constrained genes are differentially expressed.

We expect the high false negative rate of EvoGeneX in predicting adaptive genes to be because of the high variation within the subline replicates from the negative binomial representing the noise in single cell data. Despite this, we consider EvoGeneX to be powerful tool in identifying adaptive genes, specifically because of the high false positive rate of differential expression. The high contrast between the false positive rates while having similar true positive rates supports our hypothesis that differential expression alone is not enough to identify adaptive genes.

Full results are in Supplementary files Simulation_Results_EvoGeneX.xlsx and Simulation_Results_DE.xlsx.

#### 2 Lack of constrained adaptation

Interestingly, we did not detect any examples of constrained evolution in the 24 subline single cell data. There were a small number of genes (13) for which neutral evolution was rejected in favor for constrained evolution. However, constrained evolution was further rejected in favor to adaptive evolution for all of these genes. This is in contrast to results for species evolution, where, for example, at least 40% of genes were found to be constrained in Drosophila[1]. These results could result from inherent differences in selective pressures and time scales between cancer and species evolution. Within species, there are more phenotypic constraints on function within many tissues[4]. In contrast, even necessary cellular functions, like cell growth, do not face such environmental constraints in cancer and in fact are explicitly modulated in fast-growing tumors. Secondly, it could be because of the difficulty identifying genes with constrained expression in single cell data. This hypothesis is supported by simulation studies, where only 1.63% genes with expression simulated under constrained evolution are correctly predicted to be constrained. By manually examining the simulated data, the noise introduced to mimic the noise in single cell data overwhelms the signal from the constrained signal, making the resulting expression values indiscernible from neutrally-evolved data. These two hypotheses are not mutually exclusive: it is possible that cancer undergoes less constrained evolution than species and that those signals are lost due to noise from single-cell sequencing. Further examination with bulk RNA sequencing data could reveal more constrained evolution than identified here.

#### 3 Mutated genes are not more likely to have adaptive expression

In order to evaluate if mutated genes more frequently have adaptive expression, we calculated the percentage of mutated genes identified to have adaptive expression (i.e. the number of genes that are mutated in any subline and adaptive in a specific regime out of the number of genes that are mutated) and compared it to the overall percentage of genes with adaptive expression of those evaluated. We considered the 1,359 mutated genes that are also evaluated for gene expression adaptation. Of these mutated genes, approximately 9.6%, 15%, and 6.3% have adaptive gene expression for the HA-R, high aggression and sensitive (HA-S), low aggression and sensitive (LA-S) regimes, respectively. These percentages are nearly equal to the percentage of genes with adaptive expression among the 8,363 genes evaluated for adaptivity (9%, 14.4%, and 6.7% for the HA-R, HA-S, LA-S regimes, respectively) and are not significantly different based on a chi-squared test ($p = 0.98$). If we separate the mutated genes into whether the mutations are synonymous or nonsynonymous, we get similar results. This suggests that the adaptivity of a gene's expression is influenced by factors not limited to mutation status.

# 4 dN/dS

#### 4.1 Calculations

In the context of sequence evolution, a traditional way to determine adaptivity is by calculating dN/dS, the ratio of nonsynonymous mutations to synonymous mutations. This can be done both at the gene and genome level. However, the reliability of this method is debated, particularly in cancer, where assumptions made by dN/dS, like long timescales, are violated[5]. While simulation studies have indicated that the direction (if not the magnitude) of the ratio is fairly accurate[6], studies on multiple cancer types have suggested that mutational signatures, particularly high rates of C>T mutations, can bias results[7]. Further, because of the sparsity of mutations at the gene level over short time periods, dN/dS can be difficult to estimate and uninformative[8,9]. In addition, regulation of gene expression is typically mediated by non-coding regions, which are not taken into account in dN/dS analysis. Many methods have been developed to address these difficulties of dN/dS in cancer, including dNdScv, which uses a regression method to estimate the rate of mutation for each gene[10].

In order to compare our gene expression-based method to this sequence-based method, we calculated dN/dS ratios on the bulk whole exome sequence data with a counting method as well as with dNdScv[10]. Because of the sparsity of gene-wise data, we only calculate dN/dS ratios over the entire genome.

We calculate dN/dS for all sublines and the chosen sublines of each regime separately. For the counting method, we count the number of synonymous and nonsynonymous mutations across all genes in the chosen sublines. We then normalize these values by the expected number of synonymous and nonsynonymous sites in the data (1/3 of the sites are expected to be synonymous and 2/3 of the sites are expected to be nonsynonymous) and divide the proportion of nonsynonymous sites by the proportion of synonymous sites.

We also calculate dN/dS using dNdScv[10]. Same as the counting method, we calculate the ratio on all sublines and the chosen sublines of each regime separately. We input to dNdScv only the SNV mutations that occur in at least one of the selected sublines in the regime.

#### 4.2 dN/dS Results

The dN/dS ratio across all sublines by the counting method was 1.63. For the chosen sublines of each regime, dN/dS ratios are also greater than but close to one (1.32, 1.80, and 1.46 for the chosen sublines of HA-R, HA-S, and LA-S, respectively), suggesting overall slight positive selection. This is consistent with previous literature that finds that genome wide dN/dS values in cancer are close to but slightly above 1, suggesting overall slight positive selection[10].

dNdScv reported a confidence interval with mean 1.03 across all sublines. Confidence intervals for each regime also had means close to one (0.97, 1.05, 1.04 for HA-R, HA-S, LA-S respectively), suggesting near neutral evolution. Further, the over-dispersion parameter was less than one for each regime, suggesting that dNdScv may not be suitable for the dataset[10].

Complete results can be found in Supplementary Files dNdS_by_counts_Results.xlsx and dNdScv_Results.xlsx.

### 5 List of Supplementary Tables

#### 5.1 dN/dS Results

The file "dNdS_by_counts_Results.xlsx" contains the results of the dN/dS by counts method. Results of the different regimes are on different sheets within the spreadsheet. The file "dNdScv_Results.xlsx" contains the output of dNdScv. Results of the different regimes are on different sheets within the spreadsheet.

#### 5.2 Simulated Data

Separate folders for data simulated under adaptive, constrained, and neutral evolutionary models. Each file contains the simulated expression data for 100 genes for a set combination of parameters. Files are named by the parameters used. In file names, the parameters are abbreviated as: "tr": theta ratio $\theta_1/\theta_0$, "sq": Brownian motion variance $\sigma^2$, "r": dispersion ratio $r$, and "a": pull to optimum $\alpha$. The values the parameters are set to follow the abbreviation. For example the adaptive simulation file "tr_1.1_sq_.15_r_100_a_.125.csv" means that $\theta_1/\theta_0 = 1.1$, $\sigma^2 = 0.15$, $r = 100$, and $\alpha = 0.125$.

#### 5.3 Simulated Results

The results of running EvoGeneX over the simulated data is in the file Simulation_Results_EvoGeneX.xlsx. The different simulation runs are in different sheets. The sheets are labeled as "{sim type} sim; {run type} run." For example, the sheet "Neut sim; Adpt run" contains the number of genes EvoGeneX predicts to be adaptive of genes simulated under a neutral model. Each sheet also includes summary statistics for those runs and there is a "Summary" sheet containing all of those statistics.

The results of running differential expression over the simulated data is in the file Simulation_Results_DE.xlsx. The different simulation runs are in different sheets. The sheets are labeled with their simulation model. For example the sheet "Neut sim" contains the results of running differential expression over data simulated under a neutral model. Each sheet includes summary statistics for all those runs and there is a "Summary" sheet containing all of those statistics.

#### 5.4 Subline Results

Files for the results of each regime for all genes ("Gene_Results" files). These files contain the parameter estimates, q-values (BH-FDR p-values), and log fold change results for each gene for that regime. The "Clusters" file contains the cluster each gene is assigned as well as the values used to cluster the data (see Methods Section 5.4.3). The "KEGG_ORA" file contains the results of the KEGG enrichment analysis shown in Figure 4a. It contains a sheet of the number of terms in each pathway as well as sheets for each regime and regulation direction with the specific terms and associated genes.

##### 5.5 Treatment Data

The file “Bulk_Expression.csv” contains the RPKM expression values for each tumor. The columns represent tumors labeled as “X_Y_EPG” where “X” is the column number and “Y” is the mouse ID. The file “Responder_Data.csv” contains the tumors labeled the same as the “Bulk_Expression.csv” file, their response to treatment. Responses are labeled as “1R” for responder, “2S” for stable disease, “3NR” for non-responders, and “4IgG” for control tumors. It also contains the final tumor size.

##### 5.6 Treatment results

Contains a file with the output results of DESeq2 between post-treatment responder and non-responder tumors. DESeq2 results have the following columnns: “baseMean,” “log2FoldChange,” “lfcSE,” “stat,” “pvalue,” and “padj.” “baseMean” is the mean value of the responder values. “log2FoldChange” is the $\log_2$ of the mean of the responder values / the mean of the non-responder values. “lfcSE” is the standard error of the “log2FoldChange.” “stat” is the DESeq2 test statistic. “pvalue” is the raw p-value of the test statistic, and “padj” is the Benjamini-Hochberg corrected p-value.

#### References

[1] Pal, S., Oliver, B., and Przytycka, T.M. (2023). Stochastic modeling of gene expression evolution uncovers tissue-and sex-specific properties of expression evolution in the drosophila genus. J Comput Biol, 30(1), 21–40. 10.1089/cmb.2022.0121.

[2] Soneson, C. (2014). compcoder——an r package for benchmarking differential expression methods for rna-seq data. Bioinformatics, 30(17), 2517–2518. 10.1093/bioinformatics/btu324.

[3] Zappia, L., Phipson, B., and Oshlack, A. (2017). Splatter: simulation of single-cell rna sequencing data. Genome Biol, 18(1), 1–15. 10.1186/s13059-017-1305-0.

[4] Pease, J.B., Driver, R.J., de la Cerda, D.A., Day, L.B., Lindsay, W.R., Schlinger, B.A., Schuppe, E.R., Balakrishnan, C.N., and Fuxjager, M.J. (2022). Layered evolution of gene expression in “superfast” muscles for courtship. Proc Natl Acad Sci USA, 119(14), e2119671119. 10.1073/pnas.2119671119.

[5] Williams, M.J., Sottoriva, A., and Graham, T.A. (2019). Measuring clonal evolution in cancer with genomics. Annu Rev Genomics Hum Genet, 20, 309–329. 10.1146/annurev-genom-083117-021712.

[6] Pérez-Figueroa, A. and Posada, D. (2021). Interpreting dn/ds under different selective regimes in cancer evolution. bioRxiv, pp. 2021–11. 10.1101/2021.11.30.470556.

[7] Van den Eynden, J. and Larsson, E. (2017). Mutational signatures are critical for proper estimation of purifying selection pressures in cancer somatic mutation data when using the dn/ds metric. Front Genet, 8, 74. 10.3389/fgene.2017.00074.

[8] Turajlic, S., Sottoriva, A., Graham, T., and Swanton, C. (2019). Resolving genetic heterogeneity in cancer. Nat Rev Genet, 20(7), 404–416. 10.1038/s41576-019-0114-6.

[9] Chen, B., Shi, Z., Chen, Q., Shen, X., Shibata, D., Wen, H., and Wu, C.I. (2019). Tumorigenesis as the paradigm of quasi-neutral molecular evolution. Mol Biol Evol, 36(7), 1430–1441. 10.1093/molbev/msz075.

[10] Martincorena, I., Raine, K.M., Gerstung, M., Dawson, K.J., Haase, K., Van Loo, P., Davies, H., Stratton, M.R., and Campbell, P.J. (2017). Universal patterns of selection in cancer and somatic tissues. Cell, 171(5), 1029–1041. 10.1016/j.cell.2017.09.042.
