## Supplementary Tables for "Stochastic modelling of single-cell gene expression adaptation reveals non-genomic contribution to evolution of tumor subclones": Key_Resources.docx

### KEY RESOURCES TABLE

The table highlights the reagents, genetically modified organisms and strains, cell lines, software, instrumentation, and source data **essential** to reproduce results presented in the manuscript. Depending on the nature of the study, this may include standard laboratory materials (i.e., food chow for metabolism studies, support material for catalysis studies), but the table is **not** meant to be a comprehensive list of all materials and resources used (e.g., essential chemicals such as standard solvents, SDS, sucrose, or standard culture media do not need to be listed in the table). **However, please note that items in the table must also be reported in the method details section within the context of their use.**

**ALL references cited in the key resources table must be included in the main references list.** Citations should be formatted as “Author name et al.^#^” (e.g., Smith et al.^1^), with the citation number matching that in the main references list.

Please report the information as follows:

- **REAGENT or RESOURCE:** Provide the full descriptive name of the item so that it can be identified and linked with its description in the manuscript (e.g., provide version number for software, host source for antibody, strain name). See the sample tables at the end of this document for examples of how to report reagents.
  - In the **experimental models sections** (applicable only to experimental life science studies), please include all models used in the paper and describe each line/strain as model organism: name used for strain/line in paper: genotype (e.g., Mouse: OXTR^fl/fl^: B6.129(SJL)-Oxtr^tm1.1Wsy/J^).
  - The **Biological samples section** (applicable only to experimental life science studies) should list all samples obtained in this study or from commercial sources or biological repositories.
  - **You may list a maximum of 10 oligonucleotides or RNA sequences** in the table. If there are more than 10 to report, please provide this information as a supplemental document and reference the file (e.g., See Table S1 for XX) in the key resources table.
  - **Deposited data** should include both newly deposited data from this manuscript and existing datasets that were used in the manuscript.
  - Please include software and code mentioned in the method details or data and code availability section under **software and algorithms**.
  - Any item that does not fit the existing subheadings should be added to the “other” section. **Please do not add your own subheadings.**
- **SOURCE:** Report the company, manufacturer, or individual that provided the item or where the item can be obtained (e.g., stock center or repository).
  - For materials distributed by Addgene, please cite the article describing the plasmid and include “Addgene” as part of the identifier.
  - If an item is from another lab, please include the name of the principal investigator and a citation if it has been previously published.
  - If the material is being reported for the first time in the current paper, please indicate as “this paper.”
  - For software, please provide the company name if it is commercially available or cite the paper in which it has been initially described.
- **IDENTIFIER:** Include catalog numbers (entered in the column as “Cat#” followed by the number, e.g., Cat#3879S). Where available, please include unique entities such as [RRIDs](https://www.force11.org/group/resource-identification-initiative), Model Organism Database numbers, and accession numbers preceded by database abbreviations such as PDB or CCDC). Please ensure the accuracy of the identifiers, as they are essential for generation of hyperlinks to external sources when available. For more information about data sharing policies and a list of recommended data repositories for abbreviations, please see the Cell Press [Author’s guide to data sharing](https://www.cell.com/pb-assets/journals/research/cellpress/data/RecommendRepositories.pdf).
  - For antibodies, if applicable and available, please also include the lot number or clone identity.
  - For software or data resources, please include the URL where the resource can be downloaded.
  - When listing more than one identifier for the same item, use semicolons to separate them (e.g., Cat#3879S; RRID: AB_2255011).
  - If an identifier is not available, please enter “N/A” in the column.
  - ***A NOTE ABOUT RRIDs:*** we highly recommend using RRIDs as the identifier (in particular for antibodies and organisms but also for software tools and databases). For more details on how to obtain or generate an RRID for existing or newly generated resources, please [visit the RII](https://www.force11.org/group/resource-identification-initiative) or [search for RRIDs](https://scicrunch.org/resources).

Please use the empty table that follows to organize the information under the provided subheadings and skip sections that are not relevant to your study. To add a row, place the cursor at the end of the row above where you would like to add the row, just outside the right border of the table. Then press the ENTER key to add the row. Alternatively, you can right-click on your mouse and choose Insert > Insert rows above or Insert rows below. Please delete empty rows. Each entry must be on a separate row; do not list multiple items in a single table cell. Please see the sample tables at the end of this document for relevant examples in the life and physical sciences of how reagents and instrumentation should be cited.

#### TABLE FOR AUTHOR TO COMPLETE

***Please do not add custom subheadings.*** *If you wish to make an entry that does not fall into one of the subheadings below, please contact your handling editor or add it under the “other*” subheading*.* ***Any subheadings not relevant to your study can be skipped.*** *(****NOTE:*** *references should be in numbered style, e.g., Smith et al.^1^)*

**Key resources table**

| REAGENT or RESOURCE | SOURCE | IDENTIFIER |
| --- | --- | --- |
| Antibodies | | |
| Mouse anti-mouse CTLA-4 (Clone 9D9) | Bio X Cell | Cat# BE0164 |
| Mouse IgG2b (Clone MPC-11) | Bio X Cell | Cat# BE0086 |
| Bacterial and virus strains | | |
| Biological samples |  |  |
| Chemicals, peptides, and recombinant proteins | | |
| Critical commercial assays | | |
| Deposited data | | |
| Gene expression of mouse melanoma after anti-CTLA-4 treatment | This paper | GEO: GSE255484 |
| Experimental models: Cell lines | | |
| B2905 mouse melanoma cell line | Dr. Glenn Merlino  (Also available in ATCC) | B2905  (ATCC cat# CRL-3476) |
| Clonal subline of B2905, C1-C24 | Dr. Glenn Merlino | B2905 C1 to C24 |
| Experimental models: Organisms/strains | | |
| C57BL/6 mice | Charles River | C57BL/6 |
| Oligonucleotides | | |
| Recombinant DNA | | |
| Software and algorithms | | |
| Python version 3 | Python Software Foundation | [https://www.python.org](https://www.python.org/) |
| R version 4 | The R Foundation | https://www.r-project.org/ |
| EvoGeneX | Pal et al. | https://github.com/ncbi/EvoGeneX |
| clusterProfiler | Wu et al. | https://bioconductor.org/packages/release/bioc/html/clusterProfiler.html |
| dNdScv | Martincorena et al. | https://github.com/im3sanger/dndscv |
| DESeq2 | Love et al. | https://www.bioconductor.org/packages/release/bioc/html/DESeq2.html |
| Trisicell | Mehrabadi et al. | https://trisicell.readthedocs.io/ |
| Analysis code | This paper | https://github.com/ncbi/nongenetic-evolution-of-tumor-subclones |
| Other | | |

#### SAMPLE TABLES FOR AUTHOR REFERENCE

***LIFE SCIENCES***

| **REAGENT or RESOURCE** | **SOURCE** | **IDENTIFIER** |
| --- | --- | --- |
| Antibodies | | |
| Rabbit monoclonal anti-Snail | Cell Signaling Technology | Cat#3879S; RRID: AB_2255011 |
| Mouse monoclonal anti-Tubulin (clone DM1A) | Sigma-Aldrich | Cat#T9026; RRID: AB_477593 |
| Rabbit polyclonal anti-BMAL1 | This paper | N/A |
| Bacterial and virus strains | | |
| pAAV-hSyn-DIO-hM3D(Gq)-mCherry | Krashes et al.^1^ | Addgene AAV5; 44361-AAV5 |
| AAV5-EF1a-DIO-hChR2(H134R)-EYFP | Hope Center Viral Vectors Core | N/A |
| Cowpox virus Brighton Red | BEI Resources | NR-88 |
| Zika-SMGC-1, GENBANK: KX266255 | Isolated from patient (Wang et al.^2^) | N/A |
| *Staphylococcus aureus* | ATCC | ATCC 29213 |
| *Streptococcus pyogenes*: M1 serotype strain: strain SF370; M1 GAS | ATCC | ATCC 700294 |
| Biological samples | | |
| Healthy adult BA9 brain tissue | University of Maryland Brain & Tissue Bank; http://medschool.umaryland.edu/btbank/ | Cat#UMB1455 |
| Human hippocampal brain blocks | New York Brain Bank | http://nybb.hs.columbia.edu/ |
| Patient-derived xenografts (PDX) | Children's Oncology Group Cell Culture and Xenograft Repository | http://cogcell.org/ |
| Chemicals, peptides, and recombinant proteins | | |
| MK-2206 AKT inhibitor | Selleck Chemicals | S1078; CAS: 1032350-13-2 |
| SB-505124 | Sigma-Aldrich | S4696; CAS: 694433-59-5 (free base) |
| Picrotoxin | Sigma-Aldrich | P1675; CAS: 124-87-8 |
| Human TGF-β | R&D | 240-B; GenPept: P01137 |
| Activated S6K1 | Millipore | Cat#14-486 |
| GST-BMAL1 | Novus | Cat#H00000406-P01 |
| Critical commercial assays | | |
| EasyTag EXPRESS 35S Protein Labeling Kit | PerkinElmer | NEG772014MC |
| CaspaseGlo 3/7 | Promega | G8090 |
| TruSeq ChIP Sample Prep Kit | Illumina | IP-202-1012 |
| Deposited data | | |
| Raw and analyzed data | This paper | GEO: GSE63473 |
| B-RAF RBD (apo) structure | This paper | PDB: 5J17 |
| Human reference genome NCBI build 37, GRCh37 | Genome Reference Consortium | http://www.ncbi.nlm.nih.gov/projects/genome/assembly/grc/human/ |
| Nanog STILT inference | This paper; Mendeley Data | http://dx.doi.org/10.17632/wx6s4mj7s8.2 |
| Affinity-based mass spectrometry performed with 57 genes | This paper; Mendeley Data | Table S8; http://dx.doi.org/10.17632/5hvpvspw82.1 |
| Experimental models: Cell lines | | |
| Hamster: CHO cells | ATCC | CRL-11268 |
| *D. melanogaster*: Cell line S2: S2-DRSC | Laboratory of Norbert Perrimon | FlyBase: FBtc0000181 |
| Human: Passage 40 H9 ES cells | MSKCC stem cell core facility | N/A |
| Human: HUES 8 hESC line (NIH approval number NIHhESC-09-0021) | HSCI iPS Core | hES Cell Line: HUES-8 |
| Experimental models: Organisms/strains | | |
| *C. elegans*: Strain BC4011: [srl-1](http://www.wormbase.org/species/c_elegans/gene/WBGene00005636)([s2500](http://www.wormbase.org/search/variation/s2500)) II; [dpy-18](http://www.wormbase.org/species/c_elegans/gene/WBGene00001077)([e364](http://www.wormbase.org/search/variation/e364)) III; [unc-46](http://www.wormbase.org/species/c_elegans/gene/WBGene00006782)([e177](http://www.wormbase.org/search/variation/e177))[rol-3](http://www.wormbase.org/species/c_elegans/gene/WBGene00004395)([s1040](http://www.wormbase.org/search/variation/s1040)) V. | Caenorhabditis Genetics Center | WB Strain: BC4011; WormBase: WBVar00241916 |
| *D. melanogaster*: RNAi of Sxl: y[1] sc[*] v[1]; P{TRiP.HMS00609}attP2 | Bloomington Drosophila Stock Center | BDSC:34393; FlyBase: FBtp0064874 |
| *S. cerevisiae*: Strain background: W303 | ATCC | ATTC: 208353 |
| Mouse: R6/2: B6CBA-Tg(HDexon1)62Gpb/3J | The Jackson Laboratory | JAX: 006494 |
| Mouse: OXTRfl/fl: B6.129(SJL)-Oxtr^tm1.1Wsy^/J | The Jackson Laboratory | RRID: IMSR_JAX:008471 |
| Zebrafish: Tg(Shha:GFP)t10: t10Tg | Neumann and Nuesslein-Volhard^3^ | ZFIN: ZDB-GENO-060207-1 |
| *Arabidopsis*: 35S::PIF4-YFP, BZR1-CFP | Wang et al.^4^ | N/A |
| *Arabidopsis*: JYB1021.2: pS24(AT5G58010)::cS24:GFP(-G):NOS #1 | NASC | NASC ID: N70450 |
| Oligonucleotides | | |
| siRNA targeting sequence: PIP5K I alpha #1: ACACAGUACUCAGUUGAUA | This paper | N/A |
| Primers for XX, see Table SX | This paper | N/A |
| Primer: GFP/YFP/CFP Forward: GCACGACTTCTTCAAGTCCGCCATGCC | This paper | N/A |
| Morpholino: MO-pax2a GGTCTGCTTTGCAGTGAATATCCAT | Gene Tools | ZFIN: ZDB-MRPHLNO-061106-5 |
| ACTB (hs01060665_g1) | Life Technologies | Cat#4331182 |
| RNA sequence: hnRNPA1_ligand: UAGGGACUUAGGGUUCUCUCUAGGGACUUAGGGUUCUCUCUAGGGA | This paper | N/A |
| Recombinant DNA | | |
| pLVX-Tight-Puro (TetOn) | Clonetech | Cat#632162 |
| Plasmid: GFP-Nito | This paper | N/A |
| cDNA GH111110 | Drosophila Genomics Resource Center | DGRC:5666; FlyBase:FBcl0130415 |
| AAV2/1-hsyn-GCaMP6- WPRE | Chen et al.^5^ | N/A |
| Mouse raptor: pLKO mouse shRNA 1 raptor | Thoreen et al.^6^ | Addgene Plasmid #21339 |
| Software and algorithms | | |
| ImageJ | Schneider et al.^7^ | https://imagej.nih.gov/ij/ |
| Bowtie2 | Langmead and Salzberg^8^ | http://bowtie-bio.sourceforge.net/bowtie2/index.shtml |
| Samtools | Li et al.^9^ | http://samtools.sourceforge.net/ |
| Weighted Maximal Information Component Analysis v0.9 | Rau et al.^10^ | https://github.com/ChristophRau/wMICA |
| ICS algorithm | This paper; Mendeley Data | http://dx.doi.org/10.17632/5hvpvspw82.1 |
| Other | | |
| Sequence data, analyses, and resources related to the ultra-deep sequencing of the AML31 tumor, relapse, and matched normal | This paper | http://aml31.genome.wustl.edu |
| Resource website for the AML31 publication | This paper | https://github.com/chrisamiller/aml31SuppSite |

***PHYSICAL SCIENCES***

| **REAGENT or RESOURCE** | **SOURCE** | **IDENTIFIER** |
| --- | --- | --- |
| Chemicals, peptides, and recombinant proteins | | |
| QD605 streptavidin conjugated quantum dot | Thermo Fisher Scientific | Cat#Q10101MP |
| Platinum black | Sigma-Aldrich | Cat#205915 |
| Sodium formate BioUltra, ≥99.0% (NT) | Sigma-Aldrich | Cat#71359 |
| Chloramphenicol | Sigma-Aldrich | Cat#C0378 |
| Carbon dioxide (^13^C, 99%) (<2% ^18^O) | Cambridge Isotope Laboratories | CLM-185-5 |
| Poly(vinylidene fluoride-co-hexafluoropropylene) | Sigma-Aldrich | 427179 |
| PTFE Hydrophilic Membrane Filters, 0.22 μm, 90 mm | Scientificfilters.com/Tisch Scientific | SF13842 |
| Critical commercial assays | | |
| Folic Acid (FA) ELISA kit | Alpha Diagnostic International | Cat# 0365-0B9 |
| TMT10plex Isobaric Label Reagent Set | Thermo Fisher | A37725 |
| Surface Plasmon Resonance CM5 kit | GE Healthcare | Cat#29104988 |
| NanoBRET Target Engagement K-5 kit | Promega | Cat#N2500 |
| Deposited data | | |
| B-RAF RBD (apo) structure | This paper | PDB: 5J17 |
| Structure of compound 5 | This paper; Cambridge Crystallographic Data Center | CCDC: 2016466 |
| Code for constraints-based modeling and analysis of autotrophic *E. coli* | This paper | https://gitlab.com/elad.noor/sloppy/tree/master/rubisco |
| Software and algorithms | | |
| Gaussian09 | Frish et al.^1^ | https://gaussian.com |
| Python version 2.7 | Python Software Foundation | [https://www.python.org](https://www.python.org/) |
| ChemDraw Professional 18.0 | PerkinElmer | <https://www.perkinelmer.com/category/chemdraw> |
| Weighted Maximal Information Component Analysis v0.9 | Rau et al.^2^ | https://github.com/ChristophRau/wMICA |
| Other | | |
| DASGIP MX4/4 Gas Mixing Module for 4 Vessels with a Mass Flow Controller | Eppendorf | Cat#76DGMX44 |
| Agilent 1200 series HPLC | Agilent Technologies | https://www.agilent.com/en/products/liquid-chromatography |
| PHI Quantera II XPS | ULVAC-PHI, Inc. | https://www.ulvac-phi.com/en/products/xps/phi-quantera-ii/ |
